# Host aging induces a senescent-like phenotype in neutrophils and altered transcriptional responses to *Streptococcus pneumoniae*

**DOI:** 10.1101/2025.07.28.667288

**Authors:** Michael C. Battaglia, Manmeet Bhalla, Brandon Marzullo, Anagha Betadpur, Alexsandra P. Lenhard, Rania Hassan Mohamed, Murat C. Kalem, Lauren R. Heinzinger, Pathricia A. Leus, Lee Ann Garrett-Sinha, Joan Mecsas, Anna Blumental-Perry, Elsa N. Bou Ghanem

**Author notes:** Corresponding Author: Elsa N. Bou Ghanem, 955 Main Street, Buffalo, NY, 14203.

## Abstract

Aging drives increased susceptibility to respiratory infections by *Streptococcus pneumoniae* (pneumococci). Polymorphonuclear leukocytes (PMNs) are among the first responders in the lung following pneumococcal infection and are required for bacterial clearance. However, PMN antimicrobial function declines with age. To identify mechanisms underlying this decline, we performed RNA sequencing on PMNs in the lungs of young and old mice following pulmonary infection with *S. pneumoniae*. We observed significant transcriptomic differences across host age. Transcriptional analysis followed by functional validation revealed that in infected mice, PMNs from aged hosts failed to upregulate several effector activities including glycolysis and subsequent mitochondrial reactive oxygen species (ROS) production, which are necessary for bacterial killing by PMNs. Analysis of potential transcription factors controlling these changes indicated differential regulation by E2f2 in aged mice, which was linked to lower PMN differentiation resulting in more immature PMNs in the lungs of aged mice compared to young controls. Conversely, PMNs in aged mice displayed a higher senescence-associated secretory phenotype (SASP) score and upregulated pathways involved in cellular senescence. Follow-up functional characterization found that in uninfected hosts, PMNs in aged mice expressed higher levels of SASP factors IL-10, TNFα, and ROS, had lower incidence of apoptosis, and had a higher proportion of cells positive for senescence-associated β-galactosidase, features of a senescent-like phenotype. In conclusion, host aging is associated with altered PMN phenotypes, including a shift toward senescent-like energy-deficient cells, which may contribute to impaired host defense and represent potential targets for improved interventions against infection in older adults.

## Introduction

*Streptococcus pneumoniae (Sp)* is a Gram-positive bacterial colonizer of the nasopharynx that can disseminate to cause severe disease (Narciso, Dookie, Nannapaneni, Normark, & Henriques-Normark, 2024). These infections are most prevalent in age extremes, with people above 65 having higher incidence and mortality (Active Bacterial Core Surveillance Report, Emerging Infections Program Network, Streptococcus pneumoniae, 2022). As the number of older adults is projected to grow in the next 35 years from ∼56 to 95 million (Vespa, Medina, & Armstrong, 2020), understanding the mechanisms that underlie this susceptibility is vital to improving therapies and outcomes in this rapidly growing demographic.

Among the first immune responders to pulmonary infections by *Sp* are neutrophils (known as polymorphonuclear leukocytes [PMNs]). Circulating PMNs are rapidly recruited to the lungs and during infection new PMNs are generated in the bone marrow (BM) in a process called emergency granulopoiesis resulting in different PMN populations responding to the infection over time (Manz & Boettcher, 2014). PMN are vital for host defense against *Sp* as their depletion results in impaired bacterial clearance and host survival (Bou Ghanem et al., 2015; Garvy & Harmsen, 1996; McNamee & Harmsen, 2006). However, persistent PMN activity in the context of pneumonia can lead to increased bacterial numbers, tissue damage, and bacterial dissemination (Bou Ghanem et al., 2015; Taenaka et al., 2024).

PMN responses are altered with age (Bhalla et al., 2020; Fortin, Larbi, Dupuis, Lesur, & Fulop, 2007; Simell et al., 2011; Simmons et al., 2024). In the context of pneumococcal pneumonia, aged hosts have delayed initial beneficial recruitment of pulmonary PMNs with over-exuberant detrimental influx later in infection (Simmons et al., 2024) and their PMNs are defective in bacterial killing (Bhalla et al., 2020). Furthermore, adoptive transfer of PMNs from young mice reversed the susceptibility of aged mice to pneumococcal pneumonia (Bhalla et al., 2020) . This emphasizes the importance of PMNs, however, the underlying cause of their altered responses with age is not fully understood.

Immune cell dysfunction can arise from many sources including the aging process (K.-A. Lee, Flores, Jang, Saathoff, & Robbins, 2022). Aging is associated with molecular, cellular, and systemic hallmarks (Lopez-Otin, Blasco, Partridge, Serrano, & Kroemer, 2013) that affect genomic stability, metabolism, and cellular communication. These impair the ability of the host to fight infection (Ruiz et al., 2017; Stupka, Mortensen, Anzueto, & Restrepo, 2009). This impairment is in part driven by inflammaging, the low-grade chronic inflammation (Krone, van de Groep, Trzcinski, Sanders, & Bogaert, 2014) that accompanies aging and blunts the ability of immune cells to acutely respond to infection. An underpinning cause of inflammaging is cellular senescence, which is defined as a state of stable cell cycle arrest in response to stressors whereby cells remain viable and metabolically active but lose the ability to divide, become resistant to apoptosis, and develop altered responses including a senescence-associated secretory phenotype (SASP) characterized by the production of inflammatory mediators (Hodes et al., 2016). Cellular senescence plays both beneficial and detrimental roles (Gonzalez-Gualda, Baker, Fruk, & Munoz-Espin, 2021; Herranz & Gil, 2018; Hodes et al., 2016; Kuehnemann & Wiley, 2024) however, senescent cells accumulate with age and are implicated in several aging-associated diseases (Chaib, Tchkonia, & Kirkland, 2022; Gonzalez-Gualda et al., 2021; Herranz & Gil, 2018; Hodes et al., 2016).

Cellular senescence is well characterized in mitotic cells (Liu et al., 2023), but postmitotic cells can also enter a state of senescence and acquire SASP phenotype (Sapieha & Mallette, 2018; Zhao et al., 2024). In immune cells, this process is well described in T cells (Liu et al., 2023). Due to being short-lived and terminally differentiated, an age associated senescence-phenotype has not been explored in PMNs. PMNs are abundant with around 10^11^ cells produced in the BM daily (Simmons, Bhalla, Herring, Tchalla, & Bou Ghanem, 2021) that exit to the circulation as post-mitotic and terminally differentiated cells. Under homeostasis PMNs are short lived with a half-life of hours, but upon inflammation, they are recruited into tissues where they receive anti-apoptotic signals and can live up to days (Ovadia, Özcan, & Hidalgo, 2023). Given their sheer number, continuous production, and potential for extended lifespan in tissues, it is possible PMNs play a role in senescence. One study found that PMNs drive DNA damage, telomere dysfunction and senescence in hepatocytes via ROS production (Lagnado et al., 2021). PMNs showing a senescent-like phenotype were recently described in tumors (Rys & Calcinotto, 2024). In prostate cancer, apolipoprotein E (APOE) induces a senescent-like phenotype in PMNs characterized by elevated DNA damage markers, elevated ROS, reduced apoptosis, and immunosuppressive activity (Bancaro et al., 2023). In breast cancer, tumor infiltrating PMNs exhibited a senescence gene signature and elevated senescence-inducing exosomes (Ou et al., 2022). These data suggest a microenvironment-dependent induction of senescence in PMNs. However, there are no studies exploring the effect of host aging on senescence of PMNs. In this study, we assessed age-driven changes in PMN responses in a pulmonary infection model and asked if senescent-like PMNs arise because of host aging.

## Results

### PMNs display significantly different transcriptomes across host age

While PMNs were thought of as transcriptionally quiescent, findings in the past decade showed significant changes in transcription following stimulation (Montaldo et al., 2022) and tissue infiltration (Giacalone, Margaroli, Mall, & Tirouvanziam, 2020; Sumagin, 2021). To test if there are age-associated transcriptomic changes in PMNs, we performed RNA-seq on PMNs isolated from young and old mice that were pulmonary challenged with *Sp*. A highly enriched population of lung PMNs was obtained 12 and 24 hours post infection (HPI) **(SF1A-B)**. Differential gene expression analysis was conducted following bulk RNA sequencing (**Fig 1A)**. Three mice were analyzed at each timepoint. One young mouse at 24HPI was removed due to failing a quality control step. Differentially expressed genes (DEGs), defined by an absolute log2 fold-change expression of ≥.75 and a p-adj value <.05, were identified. PCA analysis showed a similar transcriptomic profile in PMNs from young and old mice 12HPI, that diverges by 24HPI **(Fig 1B)**. This coincides with a shift in PMN phenotype from protective to detrimental in the context of *Sp* infection (Bou Ghanem et al., 2015). Assessment of overlapping gene signatures indicated a conserved shift of 542 upregulated and 1,108 downregulated DEGs that occurs from 12 to 24HPI **(Fig 1C)**. However, responses were highly divergent across age with nearly 70% of DEGs identified being unique to young or old mice.

**Figure 1.**
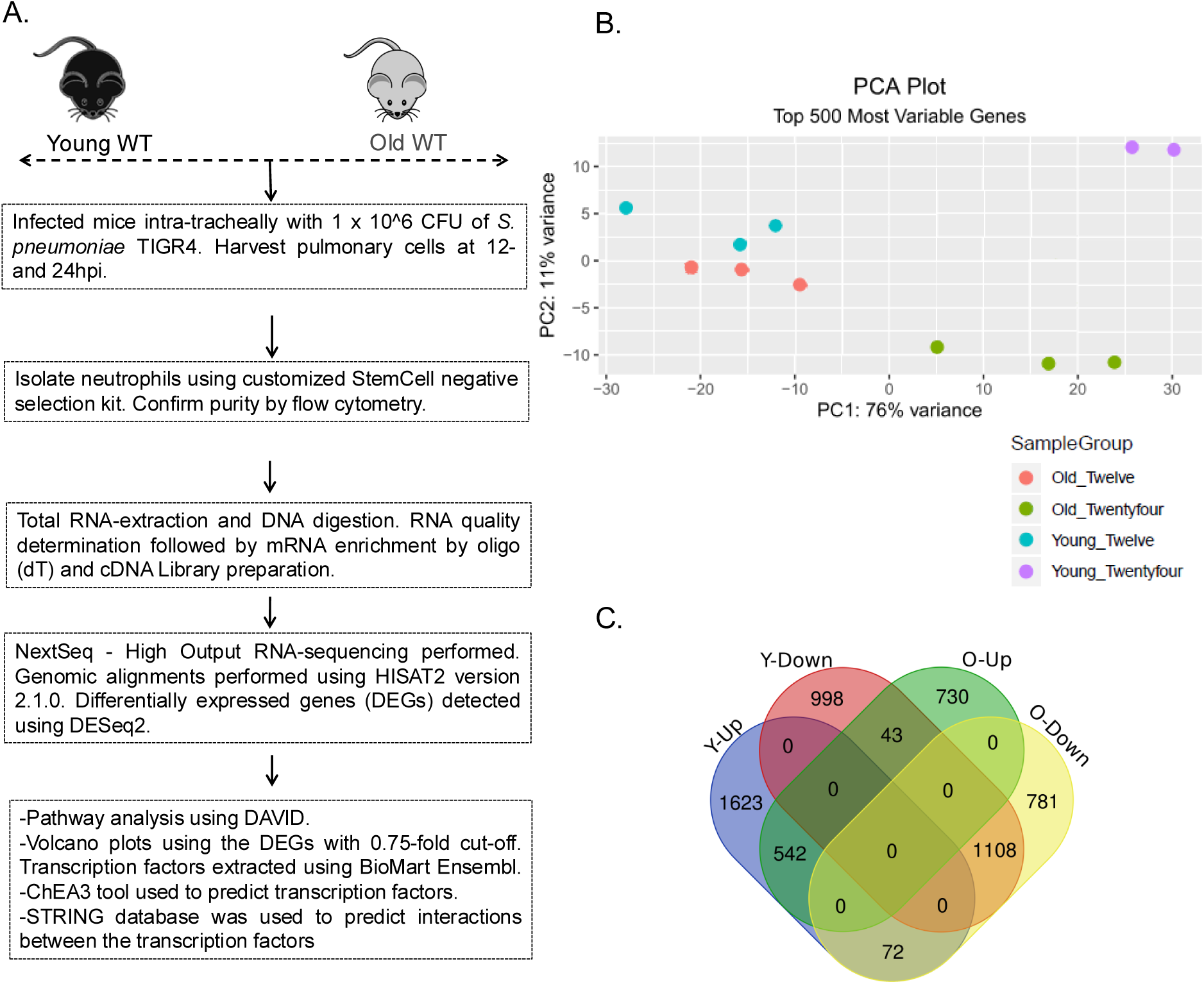
Experimental Design. Diagram of experimental layout (A). Principal component analysis (PCA) plot showing variance in expression of the top 500 variable genes in *S. pneumoniae* challenged mice (B). Venn diagram indicating overlap in DEGs identified as upregulated or downregulated in young and old mice from 12 to 24HPI (C).

We first assessed gene expression changes over the course of infection in young mice. This revealed significant up and downregulation of many genes **(SF2A)**. DAVID analysis on the DEGs identified in young mice recapitulated previous findings of upregulation of immune response pathways **(SF2B-C)**. This included enrichment of inflammatory response terms involving NF-κB, IFNγ, and TNFα, conserved proinflammatory pathways important for clearance of *Sp* by PMNs (Hackert et al., 2023; Jones, Simms, Lupa, Kogan, & Mizgerd, 2005; Khoyratty et al., 2021).

Analysis of the DEGs in PMNs from old mice revealed broadly similar overall changes in the transcriptome across time as compared to young controls **(SF3A-C)**. Of note were the enrichment of the terms “cell cycle” and “cell division” **(SF3C)**. Given that mature PMNs are terminally differentiated, this suggests that infiltrating PMNs from aged mice may be more immature and might be retaining a progenitor gene signature as influx of immature PMNs occurs during emergency granulopoiesis (Paudel, Ghimire, Jin, Jeansonne, & Jeyaseelan, 2022), which has been observed to increase in old hosts (Gullotta et al., 2023). These findings indicate an age and time-associated effect on PMN gene expression in response to *Sp* infection.

### Aging is associated with decreased PMN activation and maturation

We next wanted to study the impact of age on the gene expression at each timepoint following infection. As indicated by the PCA plot, few DEGs were identified at the 12HPI timepoint **(SF4A)** that differed between the two age groups. A significant portion of DEGs identified were part of the Igκ and heavy chain variable gene family previously reported in *Sp* infected BM PMNs from old mice (Bhalla et al., 2021). Pathway analysis of the upregulated and downregulated genes in old vs young mice at 12HPI also showed enrichment of pathways associated with the immune response in both up and downregulated genes **(Fig SF4B-C)**. Analysis of the transcriptome in young and old mice 24HPI indicated significant differential expression of many more genes **(Fig 2A)**. To determine the activation status of PMNs, we devised an Activation Score derived from normalized expression of genes annotated to the GO terms “Neutrophil Activation”, “Positive Regulation of Neutrophil Activation”, and “Negative Regulation of Neutrophil Activation”. In young mice there was significant increase in Activation Score from 12 to 24HPI that was not observed in old mice **(Fig 2B, SF5)**, where PMNs were less responsive to *Sp* infection progression. As altered effector function was previously linked to maturation status (Drifte, Dunn-Siegrist, Tissieres, & Pugin, 2013), we assessed that. Utilizing a gene list derived from sc-RNAseq (Ai, 2022), we generated a normalized expression Maturation Score. The results mimicked that of the Activation Score with young mice displaying a significant increase in PMN maturation genes during infection **(Fig 2C, SF6)**, which was not observed in old mice **(Fig 2C)**. Together, these data suggest that PMNs that influx into the lung of old mice later in infection are less mature and less capable of activation. In line with these data, pathway analysis of DEGs downregulated in old versus young mice 24HPI indicated enrichment of pathways involved in immune responses **(Fig 2D)**.

**Figure 2.**
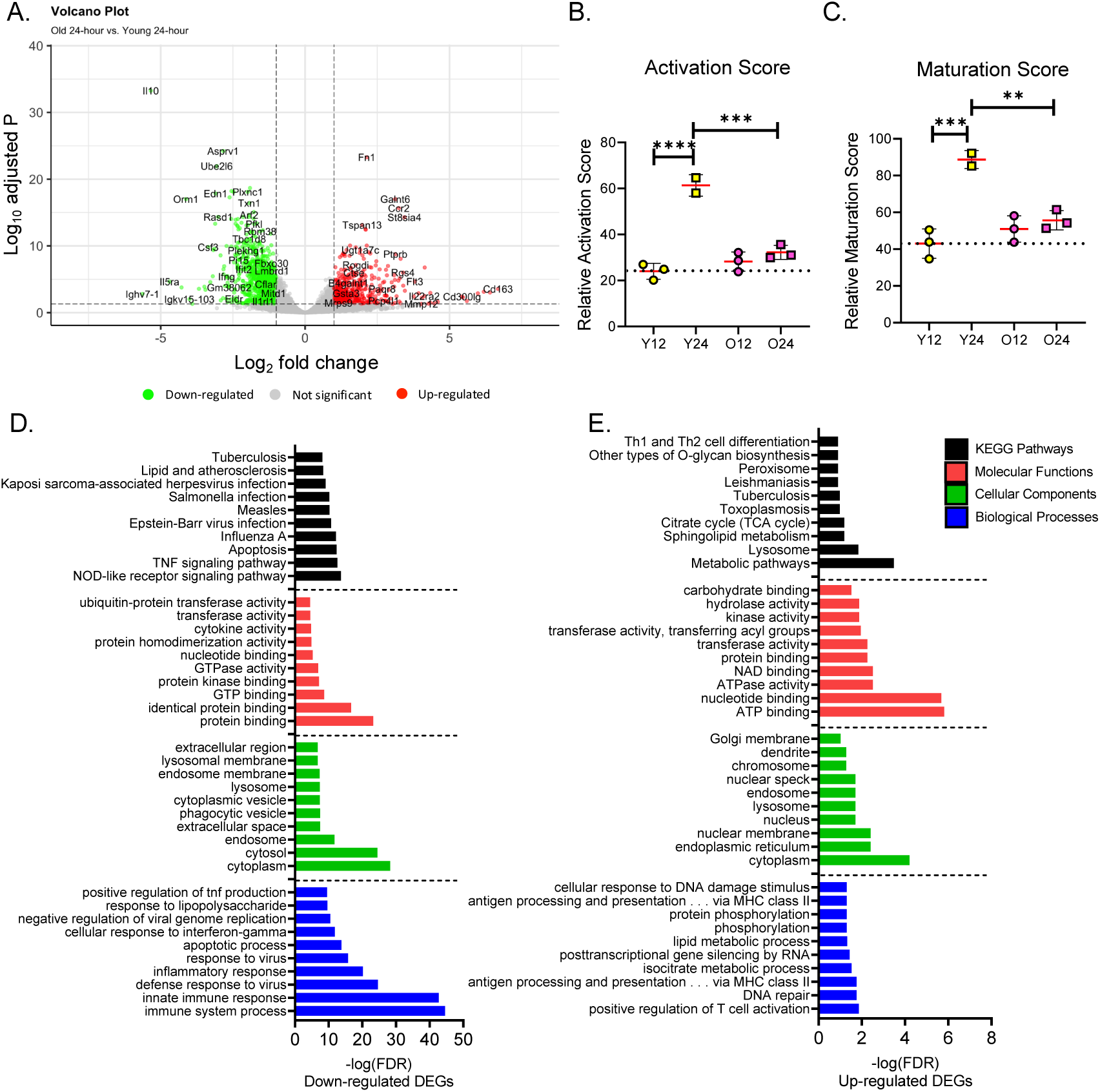
Transcriptome of lung PMNs in aged hosts is associated with reduced activation and maturation. Volcano plot of DEGs identified between old and young mice at 24HPI (A). Activation (B) and maturation (C) scores for lung PMNs from young and old mice at 12 and 24HPI. Top 10 results of DAVID analysis for terms enriched in DEGs identified as downregulated (D) and upregulated (E) in old vs young mice at 24HPI.

To assess if aging alters expression of PMN effector function related genes, we focused on genes involved in granule production, phagocytosis, and reactive oxygen species (ROS) production. PMN granules are categorized as primary, secondary, and tertiary defined by their contents (Othman, Sekheri, & Filep, 2022). Primary granules develop early in granulopoiesis (Lehman & Segal, 2020), and carry out antibacterial function while contributing to lung pathology through damage of epithelial tissues (Dickerhof et al., 2020; Haegens et al., 2009; Voynow & Shinbashi, 2021). PMNs from old hosts exhibited elevated expression of many primary granule components including Elane and Mpo at 24HPI compared to young host **(SF7)**. This corroborates previous results indicating increased neutrophil elastase activity in older donors (Bou Ghanem et al., 2017) and suggests that PMNs in old mice may be more damaging to tissues (Xu et al., 2024). Conversely, PMNs from young hosts exhibited an infection dependent elevated expression of secondary **(SF8)** and tertiary **(SF9)** granules. These granules, which develop later in PMN maturation, have roles in pathogen clearance and tissue remodeling (Gigon, Yousefi, Karaulov, & Simon, 2021). As granule development and content expression are heavily linked to PMN maturation, these data further suggest immaturity of lung PMNs in old mice.

Phagocytosis is required for pneumococcal killing by PMNs (Siwapornchai et al., 2020), while Nox-mediated ROS production has a limited role in PMN-mediated killing of *Sp* (Herring et al., 2022) but is important for killing of other pathogens (Rada, Geiszt, Káldi, Timár, & Ligeti, 2004). PMNs from young hosts exhibited a near uniform increase in response to infection over time in expression of genes involved in phagocytosis, phagosome maturation, and Nox **(SF10-11)**. Conversely, PMNs from the old host exhibited little to no change in expression of genes related to phagocytosis and Nox upon progression of the infection from 12 to 24HPI **(SF10-11)**. These findings confirm previous reports of aberrant effector functions in PMNs from old hosts, validating the dataset (Simmons et al., 2021).

### *E2f2* gene signature is enriched in PMNs from old mice

While pathway analysis provides clues as to the altered function and maturity of PMNs, it does not provide mechanisms that underlie these changes. To assess if there are altered transcription factor (TF) networks in PMNs that impact the gene expression, we utilized the tool ChEA3, which predicts TF enrichment based on integration of Encode ChIP and co-expression data (Keenan et al., 2019). Through analysis of the DEGs up and downregulated in both age groups, a total of 98 TFs (24 unique to young, 34 unique to old, 40 overlapping) were identified as significant (TopRanked Score<.01) **(Fig 3A)**, with many previously identified as controlling PMN development and function (Xie et al., 2020). Pathway analysis of the identified TFs enriched in young mice support previous data suggesting a robust immune response by pulmonary infiltrating PMNs **(SF12 A, C)**, with significant enrichment of terms such as “response to cytokine” and “innate immune response”.

**Figure 3.**
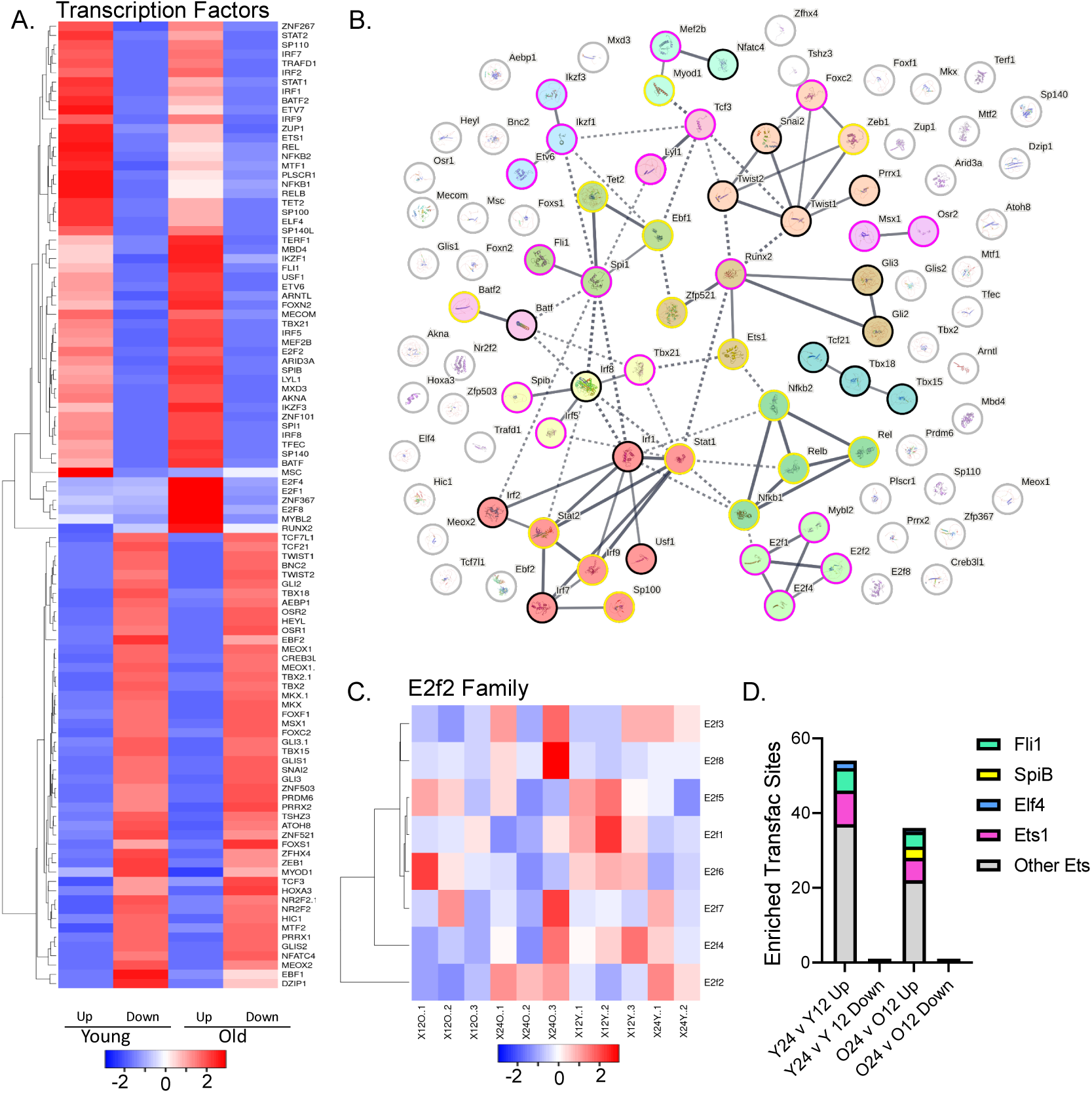
Alterations in transcriptome is associated with differential gene signature for E2f2 and Ets1. Heatmap showing -log(TopRanked Score) from significant results of ChEA3 analysis of up- or downregulated DEGs in lung PMNs of young and old mice 24HPI vs 12HPI (A). STRING analysis displaying high confidence interactions for transcription factors identified by ChEA3 in young mice (yellow halo), old mice (pink halo), or both (black halo) clustered using MCL analysis (B). Heatmap of E2f family transcription factors expression (C). TRANSFAC analysis of upregulated and downregulated DEGs in young and old mice 24HPI vs 12HPI highlighting Ets transcription factors (D).

To identify the networks regulated by the TFs identified by ChEA3, we used STRING software (Szklarczyk et al., 2023), a tool that shows physical and regulatory interaction networks. STRING analysis identified 13 functional clusters of TFs **(Fig 3B Table S1)** including enrichment of factors in upregulated DEGs of old mice controlling cell cycle progression (cluster 6), which was primarily composed of members of the E2F family. The E2F TF family is composed of eight members involved in cell cycle, differentiation, DNA damage responses, and apoptosis (Morgunova et al., 2015), which are often affected in senescence (Liu et al., 2023). Gene expression analysis of the E2F family revealed increase in E2f2 expression in response to *Sp* infection over time in PMNs from both young and old mice **(Fig 3C)**.

The ChEA3 analysis also revealed enrichment of Ets family TFs **(Fig 3A-B)**, which are important for lymphocyte development and function (Sharrocks, 2001). Due to similar DNA-binding motifs, Ets1 family members often share similar targets (Wei et al., 2010). In order to isolate potentially important Ets family members, we utilized TRANSFAC (Matys et al., 2006), another TF prediction tool, whose results were cross referenced with that of ChEA3 resulting in identification of Ets1 **(Fig 3D)**. Ets1 is a TF expressed in lymphoid cells that controls their development and function (Garrett-Sinha, 2023; C. G. Lee et al., 2019; Sunshine et al., 2019), but has not been studied in PMNs.

We then wanted to study the role of Ets1 and E2f2 in PMN differentiation and effector function. As full knockouts of Ets1 (Garrett-Sinha, 2023) and E2f2 (Murga et al., 2001; Zhu et al., 2001) have broad systemic effects and primary PMNs are difficult to genetically manipulate, we used the Hoxb8 system, which allows for CRISPR/Cas9 mediated gene knockouts and generation of PMNs *in vitro* (Shannon & Hinnebusch, 2023; G. G. Wang et al., 2006). Hoxb8 derived PMNs were generated as previously described (Nguyen et al., 2020) resulting in CD11b^+^Ly6G^+^ cells **(Fig 4A)**. Loss of Ets1 had no impact on PMN maturation where Ets1KO cells displayed similar prevalence of mature CD11b^+^Ly6G^+^ cells as compared to the parental line **(Fig 4B)**. As Ets1 may play a role in activation of PMNs, similar to its role in B and T cells (Garrett-Sinha, 2013), we assessed the killing capacity of BM PMNs isolated from Ets1+/+, Ets1+/-, and Ets1-/- mice. Sera from the wild-type Ets1+/+ and Ets1-/- mice were used for opsonization to control for autoimmune mediated hypocomplementemia (Garrett-Sinha, 2023). PMNs isolated from all three genotypes exhibited similar killing regardless of the opsonin **(SF13A)**. In addition, Ets1 transcript levels did not change upon *in vitro* infection of PMNs isolated from human donors **(SF13B)**, suggesting Ets1 does not play a role in PMN function.

**Figure 4.**
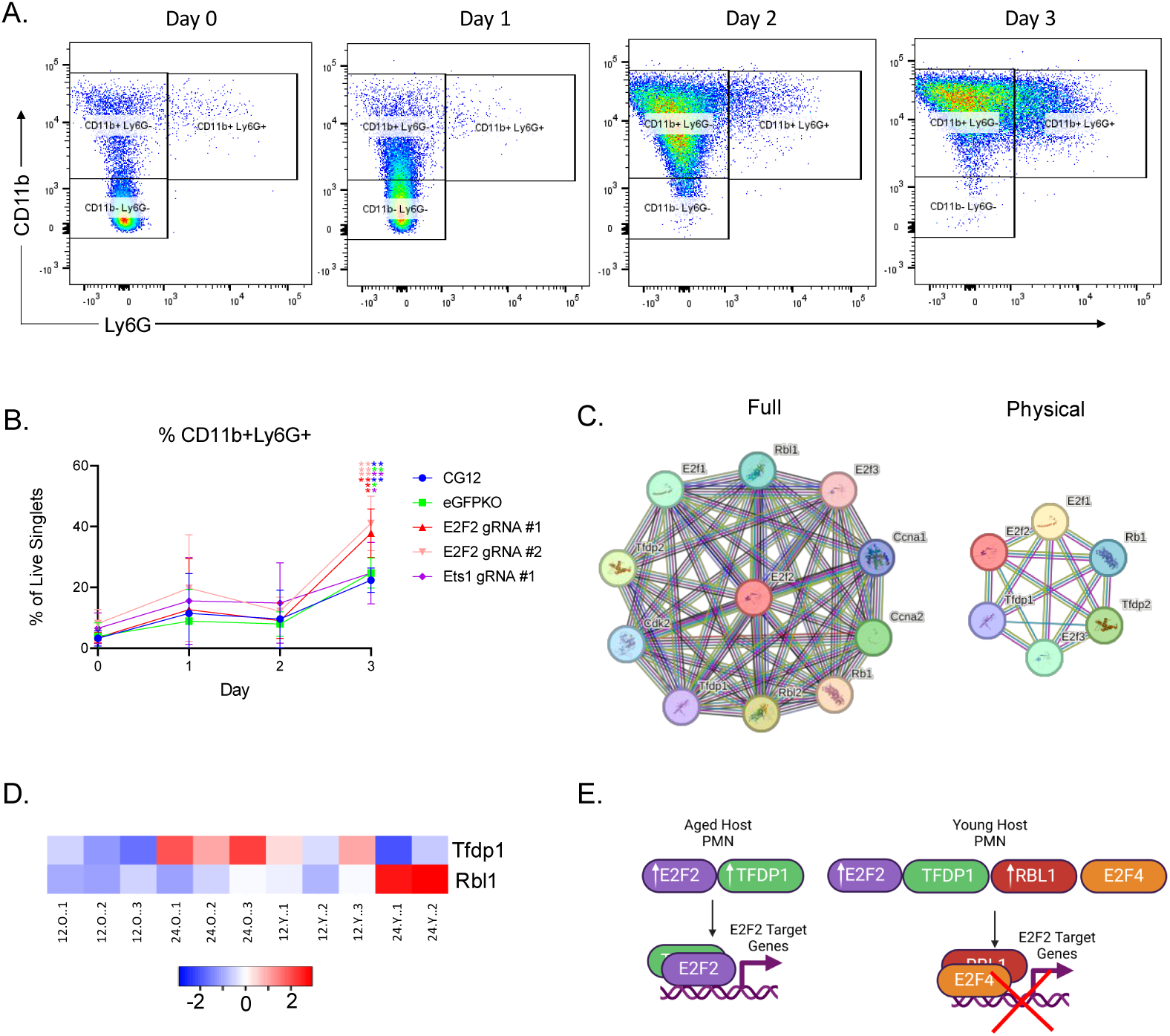
E2f2 negatively regulates PMN maturation. Representative data showing CD11b vs Ly6G expression and gating strategy for HoxB8 cells during differentiation on days 0, 1, 2, and 3 (A). Percent of CD11b+Ly6G+ cells on days 0, 1, 2, and 3 of HoxB8 differentiation (B). Data are pooled from 4 separate experiments and statistical significance was assessed using one-way ANOVA followed by Tukey’s multiple comparisons test (B). Full/physical STRING interaction networks for E2f2 (C). Heatmap showing normalized expression for Tfdp1 and Rbl1 (D). Proposed model for differential regulation of E2f2 in lung PMNs in young and old hosts showing differential regulation of E2f2 regulators (E).

Conversely, analysis of two generated Hoxb8 lines deficient for E2f2, using different gRNAs, displayed an increase in fully mature cells generated post differentiation as compared to the parental control (CG12) and the knockout system control (eGFPKO) **(Fig 4B)**. This suggests that E2F2 negatively regulates PMN maturation. As E2f2 was significantly upregulated in both young and old mice over time **(Fig 3C)**, while the downstream gene signature for E2f2 was only identified as significantly enriched in old mice **(Fig 3A-B)**, we asked if there was differential regulation of E2f2 activity via interacting partners. Using STRING, we identified high confidence interactors of E2f2 **(Fig 4C)** with two that were DEGs in our dataset. Tfdp1 is an heterodimeric partner of E2f2 that is required for DNA binding (Morgunova et al., 2015) and that was upregulated upon progression of infection in old mice **(Fig 4D)**. In contrast, in young mice we observed a significant upregulation of Rbl-1 **(Fig 4D)**. The E2f family member E2f4 exists in the cytosol where upon binding to Rbl-1, it translocates to the nucleus where it exerts repressive activities on E2f2 target genes (B.-K. Lee, Bhinge, & Iyer, 2011; Morgunova et al., 2015). The upregulation of Rbl-1 in young mice suggests that despite the upregulation of E2f2 expression, its activity is decreased resulting in maturation of PMNs. In PMNs from old mice, we observed an increase in expression of both E2f2 and its positive regulator Tfdp-1 suggesting enhanced E2f2 activity and reduced maturation of newly generated PMNs **(Fig 4E)**.

### Aging is associated with altered metabolic signatures in pulmonary PMNs

Metabolic shifts are often seen in age-associated senescence of immune cells that affect function and differentiation (Wiley & Campisi, 2021). Activated PMNs rely heavily on aerobic glycolysis, which has been shown to be important for ROS production, phagocytosis, and NETosis (Awasthi et al., 2019; Ettel & Weichhart, 2024; Toller-Kawahisa et al., 2023). Pathway analysis indicated metabolic shifts in pulmonary infiltrating PMNs with significant upregulation of genes involved in the Citric Acid Cycle (TCA) **(Fig 2E)** in old vs young hosts. In contrast, in young mice there was upregulation of nearly all genes involved in glycolysis accompanied by downregulation of nearly all TCA components during the course of infection **(SF14A-C)**. Conversely, old mice displayed the opposite trend with little to no change in expression of glycolytic components and an upregulation of TCA component genes over time **(SF14A-C)**. PMN precursors in the BM and circulating immature PMNs were reported to increase utilization of oxidative phosphorylation (Riffelmacher et al., 2017), which is immediately downstream of the TCA cycle. This again suggests that PMNs in old mice during late stage of infection are more immature relative to young controls.

To assess whether glycolysis is required for bacterial killing, PMNs from young mice were treated with 2-DoG or lonidamine, known inhibitors of glycolysis. In a dose dependent manner, we found that glycolysis inhibition reduced opsonophagocytic killing of *Sp* **(Fig 5A)**. Similarly, we found that inhibition of glycolysis by 2-DoG resulted in near complete abrogation of *S. pneumoniae* killing by human PMNs from five donors **(Fig 5B)**. These data corroborate the previous work (Fan et al., 2021), showing requirement of glycolysis for *Sp* killing.

**Figure 5.**
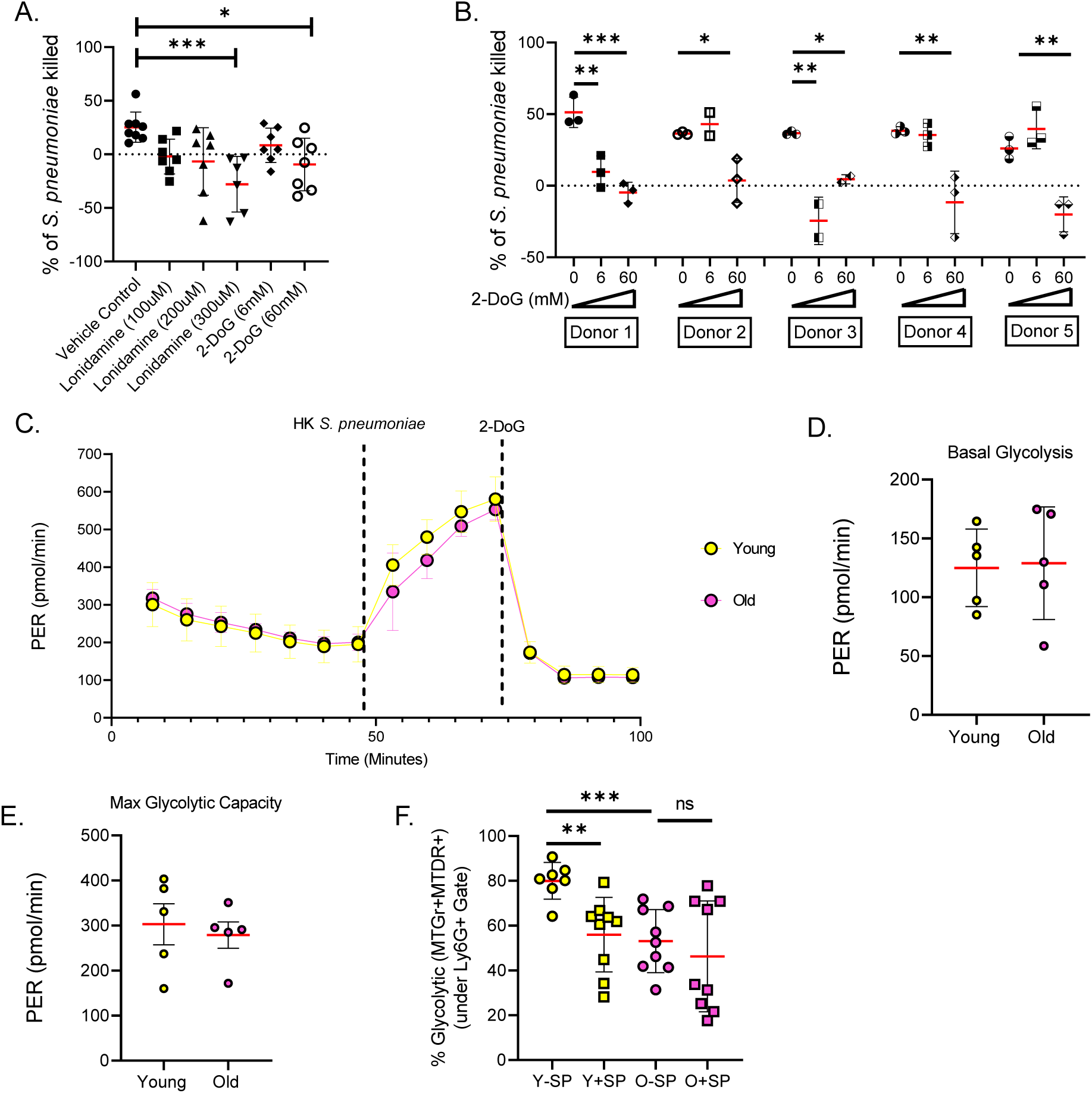
Glycolysis upregulation is required for PMN killing of *S. pneumoniae*. Percent bacterial killing using BM PMNs from young mice treated with VC, lonidamine, or 2-DoG at the listed concentrations (A). Data are pooled from 6 experiments with significance determined by one-way ANOVA followed by Tukey’s multiple comparisons test (A). Data showing technical replicates of bacterial killing using human peripheral PMNs from 5 separate donors in the presence of a VC or 2-DoG at the listed concentrations (B). Significance was determined by one-way ANOVA (B). Representative results of Glycolysis Stress Test conducted on BM PMNs from young and old mice (C). Basal glycolysis (D) and max glycolysis (E) measurements as derived from Glycolysis Stress Test. Data are pooled from 5 experiments each with significance determined by unpaired t-test (C-E). (F) Flow cytometry data showing % glycolytic PMNs in the lung of young and old mice relative at baseline and 24HPI. Data are pooled from 7-9 mice per group across three separate experiments (F). Statistical significance was determined by One-way-ANOVA followed by Tukey’s multiple comparisons test.

To determine whether the change in metabolic gene expression constituted an intrinsic shift in the metabolic capacities of PMNs, we assessed glycolytic capacity using Agilent Seahorse. For feasibility with the numbers needed for the assay, we focused on BM PMNs. As PMNs are primarily dependent on glycolysis for their energetic needs, inhibition of respiration is not sufficient to reach maximal glycolytic capacity (Grudzinska et al., 2023). We therefore used opsonized heat-killed (HK) *Sp* as a stimulus to mimic pathogen specific responses **(SF15A),** which increased glycolysis beyond that induced by respiration inhibition. We found that PMNs from young and old mice displayed similar basal glycolysis and max glycolytic capacity **(Fig 5C-E)**. As pathway analysis indicated upregulation of the TCA cycle **(Fig 2D)**, we also tested if there were differences in the respiratory capacity of PMNs. To do so, we utilized a Cell Mito Stress Test to characterize respiration **(SF15B)**. Results indicated no significant differences in basal respiration **(SF15C),** max respiration **(SF15D)**, spare capacity **(SF15E)**, and ATP production **(SF15F)**. Nevertheless, all bio-energetic parameters were trending to be lower in old mice, indicating that mitochondrial ability to produce energy slowly declines, with great variability in individual PMNs, as their aging is not homogenous/synchronized. In agreement with less efficient ATP production, the proton leak was most reduced.

To assess if the energetic state of PMNs is altered in the lungs, we used flow cytometry to assess glycolytic (MitoTracker Gr^+^ MitoTracker DR^+^) and respiratory (MitoTracker Gr^-^ MitoTracker DR^+^) PMNs in the lungs as previously described (Díaz-Basilio et al., 2024). In the absence of infection young mice displayed a significantly higher portion of glycolytic PMNs as compared to old mice **(Fig 5F)**. Upon infection, the proportion of glycolytic PMNs decreased in young mice corresponding with increase in respiratory PMNs **(SF15H)**. In old mice there was no observed change in PMN energetics following infection **(Fig 5F, SF15H)**. Together these data indicate significant changes in the metabolic pathways in PMNs in a tissue environment dependent manner.

### PMNs in old hosts display a senescence-like phenotype

Senescent cells are characterized by increased lysosomal content, cell cycle arrest, a secretory phenotype, apoptotic resistance, and changes in metabolism (Hernandez-Segura, Nehme, & Demaria, 2018). To assess if aging is associated with a senescence-like phenotype in PMNs, we examined the RNAseq dataset. Pathway analysis of DEGs upregulated in old relative to young mice at 24HPI found enrichment of pathways related to DNA damage and respiration including “DNA repair” and “Citrate cycle (TCA)” **(Fig 2E)**. Similarly, terms associated with TFs enriched in DEGs from PMNs in old mice suggest an immune response with a senescent phenotype **(SF12B-D)** with enrichment of terms such as “immune system process”, “regulation of cell cycle”, and “cellular senescence”. One hallmark of cellular senescence is the senescence-associated secretory phenotype (SASP) (Li et al., 2023). To determine if this was evident in the data, we devised a score based on the normalized expression of 22 SASP factors (Bleve et al., 2023; Chaib et al., 2022; Lagnado et al., 2021). Analysis at 12HPI indicated an increased SASP phenotype in old mice driven by significantly higher expression of factors such as IL-10 and TNFα **(Fig 6A SF16A)**. Examining pathways identified by DAVID revealed that at 12HPI there was significant enrichment for several pathways associated with a senescent-like phenotype **(Fig 6B)**.

**Figure 6.**
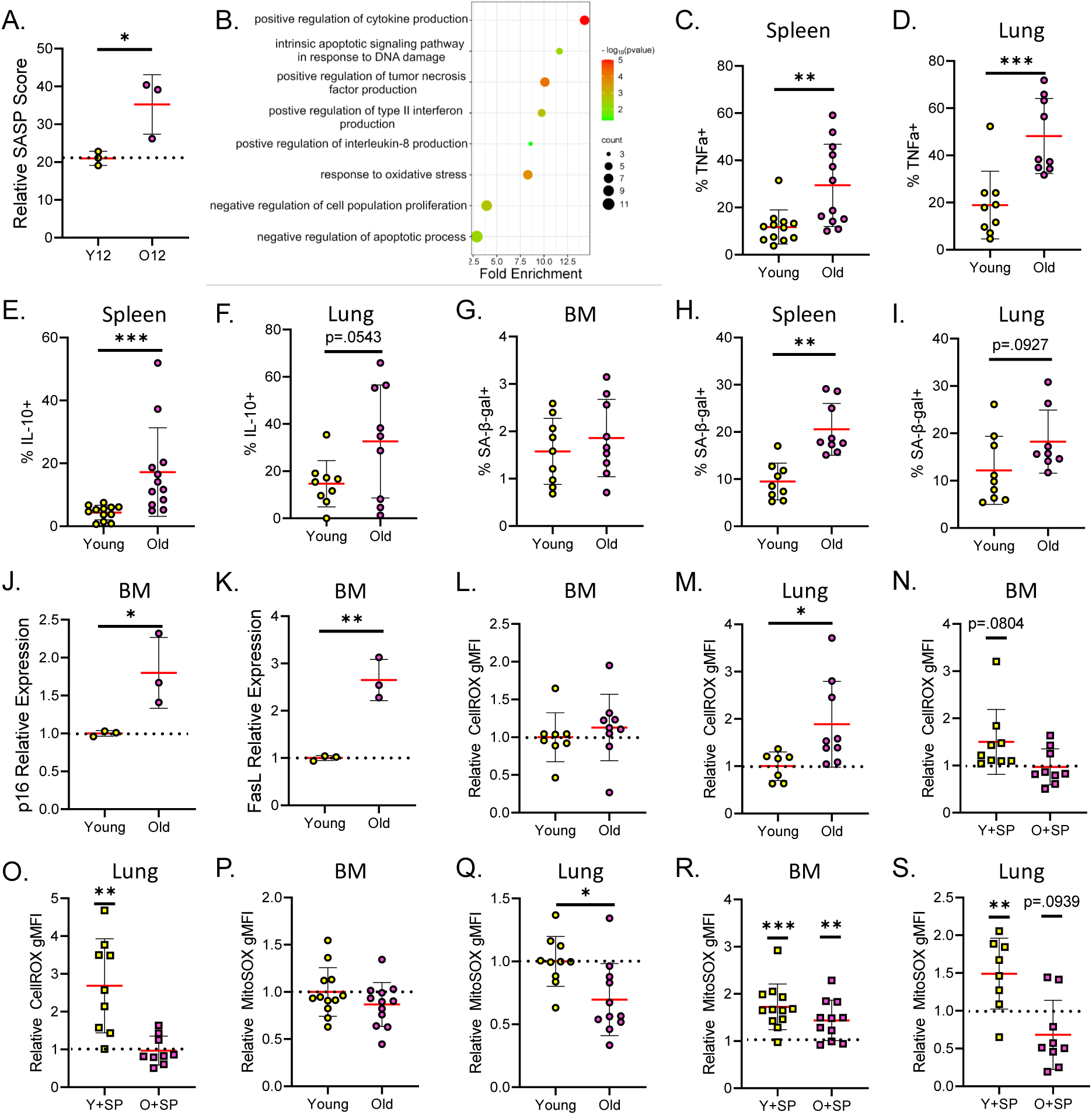
Old mice display a senescence-like phenotype in PMNs. Relative SASP Score of lung PMNs in young and old mice 12HPI with significance determined by unpaired t-test (A). Bubble plot showing curated list of GO terms enriched in PMNs in old vs young mice at 12HPI (B). Flow cytometry data showing % TNFα+ and %IL10+ PMNs in lung (C, D) and spleen (E, F) of young and old uninfected mice. Flow cytometry data showing %SA-β-gal+ PMNs in the BM (G), spleen (H), and lung (I) of young and old uninfected mice. RT-qPCR data showing relative expression of p16 (J) and FasL (K) in BM PMNs of young and old uninfected mice. Flow cytometry data showing relative gMFI for CellROX in the BM (L) and lungs (M) of young and old uninfected mice normalized to young controls. Flow cytometry data showing gMFI for CellROX in BM (N) and lung (O) and MitoSOX in the BM (P) and lung (Q) PMNs in young and old uninfected and infected mice relative to own uninfected controls. Flow cytometry data showing gMFI for MitoSOX in BM (R) and lung (S) PMNs in young and old uninfected and infected mice relative to matched uninfected controls. Data are pooled from 9 mice per group across three separate experiments (C-S). Statistical significance was determined by unpaired t-test between indicated groups (D, F, G, J, K, M, Q) or with respect to uninfected controls (L, N, O, P, R, S) or Mann-Whitney test (C, E, H, I).

To validate the age-associated senescence-like PMN phenotype, we measured several markers *in vivo* in the absence of infection. We first assessed production of SASP-related cytokines TNFα and IL-10 in PMNs of peripheral organs including lungs and spleen. Analysis of splenic PMNs in old mice indicated a significant increase in the percent of cells expressing TNFα **(Fig 6C)** and IL-10 **(Fig 6E),** which was mimicked in relative expression (**SF17A/C)**. These expression patterns were also observed in lung PMNs **(Fig 6D/F SF17B/D)**. Together these data suggest that at baseline, PMNs from old hosts display a SASP phenotype in peripheral sites.

Another feature of cellular senescence is increased activity of senescence-associated β-galactosidase (SA-β-gal) (B. Y. Lee et al., 2006). Analysis of PMNs in the absence of infection indicated no difference in SA-β-gal+ PMNs in the BM **(Fig 6G)**. However, analysis of PMNs in the spleen **(Fig 6H)** and lungs **(Fig 6I)** revealed increases in SA-β-gal+ PMNs in aged compared to young hosts. Interestingly, most SA-β-gal+ PMNs in the spleen and lung displayed a Ly6G^lo^ phenotype, associated with immature PMNs, which was more prevalent in old mice **(SF17E-F)**. This suggests that there is an age-related SA-β-gal expression primarily in immature PMNs in the periphery.

Senescence is also associated with elevated DNA damage response, reduced proliferation, and an accompanied resistance to normal apoptotic clearance (Hernandez-Segura et al., 2018; Hu et al., 2022). In the absence of infection, PMNs within the BM of old mice exhibited elevated expression of p16 **(Fig 6J)**, a cell cycle arrest protein that is upregulated in the event of DNA damage (Rodier et al., 2009), FasL **(Fig 6K)**, a programed cell death receptor ligand reported to be upregulated in many senescent cells (Lagunas-Rangel, 2023), and lower expression of Ki67 indicative of lower proliferation **(SF17G)**. This suggests that there is senescent-like phenotype emerging in PMNs in the bone-marrow niche where they are generated.

We next assessed if there was a senescent-like resistance to apoptosis (Hu et al., 2022). As expected, in young hosts, we observed upregulation of most genes involved in apoptosis that was not observed in old mice **(SF16B-C)**. To assess if there was alteration *in vivo,* we measured apoptotic PMNs in the lungs. We observed a greater proportion of old mice with low levels of apoptosis (<10%) as compared to young mice **(SF16D)**. Together, these data suggest that PMNs in old hosts experience greater apoptosis resistance in the lungs.

As our data suggested increased oxidative stress in old mice, we then measured ROS at baseline and after infection. We found that at baseline, PMNs in the lungs but not BM from old mice had increased production of ROS **(Fig 6L-M)**. Following infection, BM PMNs in young mice but not old mice displayed a trend towards increased ROS **(Fig 6N)**, which was significantly enhanced in the lung **(Fig 6O)**. This implies that lung PMNs in old hosts are experiencing greater oxidative stress even in the absence of infection and are unable to respond to acute stimuli. We next measured production of mitochondrial ROS (mitoROS), which are vital for PMN mediated clearance of *Sp* (Herring et al., 2022). We found that PMNs in old mice produced similar mitoROS in the BM and significantly lower mitoROS in the lungs as compared to young controls **(Fig 6P-Q)**. Within the BM, PMNs from young and old mice displayed similar mitoROS increases relative to their own baselines following pulmonary *Sp* infection **(Fig 6R)**. However, lung PMNs of old mice failed to upregulate mitoROS production in response to infection as observed in young mice **(Fig 6S)**. This indicates that PMNs in old hosts are unable to properly upregulate protective mitoROS following migration from the BM to the lungs. This suggests that the lung microenvironment in old mice is abrogating PMN acute responses to *Sp* infection.

PMNs display an aging phenotype characterized by loss of expression of CD62L and increase expression of CXCR4 and CD49d, in a circadian rhythm dependent manner (María Casanova-Acebes et al., 2013) that have altered function (Ovadia et al., 2023). To test if the observed increase in senescent-like PMNs is indicative of increased aged PMNs, we tested for their presence by flow cytometry. Analysis indicated that at baseline, old mice displayed similar presence of aged (CD49d+CXCR4+CD62L-) PMNs in the BM **(SF17H)** but decreased prevalence in the lung **(SF17I**), which fits our observation that peripheral PMNs in old mice display a more immature phenotype **(Fig 2C)**.

All together, these data suggest that PMNs in aged hosts acquire phenotypes like that of senescent cells as early as in the BM. However, many facets of senescence-like phenotype are not evident until the cells migrate to the periphery. In the periphery PMNs from old mice displayed altered secretory phenotype, apoptosis, and ROS production at baseline and in response to *Sp* infection, indicative of adoption of a senescence-like phenotype that is distinct from aged PMNs described previously.

## Discussion

Here we examined the effect of host aging on transcriptional changes in PMNs during *Sp* pulmonary infection. In young mice our results are consistent with previous studies finding significant upregulation of effector function related genes following activation of PMNs *in vitro* and *in vivo* (Gomez, Dang, Martin, & Doerschuk, 2016; Gour et al., 2024; Khoyratty et al., 2021; Lu et al., 2021; Matarazzo et al., 2024; Minhas et al., 2020). However, analysis of the transcriptome in old hosts revealed a significantly different response characterized by reduced effector functions, altered metabolism and maturation, and a senescent-like phenotype. *In vivo* functional confirmation of these changes revealed tissue specific phenotypes, where several of the age-driven alterations were observed in the lungs but not in the BM. This suggests that it is in the periphery that many facets of PMN dysfunction are acquired during host aging. It further suggests that heterogeneity in PMN phenotype arises in part in response to tissue instruction (M. Casanova-Acebes et al., 2018; Ganesh & Joshi, 2023). These changes may be due to altered lung microenvironment during aging mediated by elevated cytokine or damage related signaling molecules (Leblanc et al., 2024). Whether these changes are restricted to a *bona fide* subset that arises in the BM (Xie et al., 2020) and acquire phenotypes in the periphery is an open question.

In examining PMN metabolism in infected and uninfected states we found host aging results in elevated respiration in PMNs. As PMNs have been primarily described as glycolytic (Ettel & Weichhart, 2024; Fan et al., 2021; Leblanc, Bourgoin, Poubelle, Tessier, & Pelletier, 2024) this may be indicative of age-related energy deficit, where PMNs from old hosts are unable to maintain their energetic requirements through glycolysis, despite similar concentrations of glucose being available. We hypothesize that this prompts compensatory elevated respiration to generated energy, which is reflected in changes in gene expression profile during infection progression. Previous studies have found that in many cases aging correlates with decreased mitochondrial function (Gómez & Hagen, 2012). As our data suggests that mitochondria in lung PMNs in young but not old mice become more respiratory in response to infection, it is possible that PMNs from old mice have dysfunctional mitochondria that are unable to acutely respond and boost energy production when challenged. This deficit in energy production may in part account for PMN reduced effector capacity. Many of these metabolic shifts have been characterized in B and T cells during aging (Frasca, Diaz, Romero, Garcia, & Blomberg, 2020; Liu et al., 2023). Significant changes were observed in T cell metabolism following development of age related senescence including reduced cellular respiration (Ron-Harel et al., 2018) and ATP production (Fang et al., 2016). While there is a classical reliance on glycolysis in senescent cells (Liu et al., 2023), it was observed that glycolysis can be reduced in T cells in aging hosts (Ron-Harel et al., 2018). Our data suggest that this is also true for PMNs, which are experiencing a senescent-like state in the lung microenvironment, have lower basal glycolysis and are unable to acutely upregulate cellular respiration in response to *Sp* infection. Together these studies suggest that in aging there may be a shared dysregulation of carbon metabolism in multiple immune cell types in response to exogenous activation.

We observed age-driven decrease in PMN maturation that was linked to E2f2 expression. While the role for E2f family members in PMN development is not well studied, studies using *in vitro* granulocyte differentiation indicated that E2F target genes were downregulated in transitioning from promyelocytes to myelocytes, suggesting that the E2f2 signature in old hosts may be associated with immature PMNs (Theilgaard-Mönch et al., 2005). Similarly, studies in COVID-19 indicated patients with an enriched E2F gene signature in leukocytes displayed increased immature PMNs that correlated with worse outcomes (Lam et al., 2023). In addition, previous studies have indicated that E2f2 is associated with a SASP phenotype in synovial fibroblasts (Bao & Hu, 2018; S. Wang, Wang, Wu, Sun, & Pan, 2018; R. Zhang, Wang, Pan, & Han, 2018). Together these data suggest that E2f2 negatively controls maturation of functional PMNs, and that maintenance of its activity correlates with a senescence-like phenotype.

Lastly, we observed a senescent-like phenotype in PMNs that emerges primarily in the periphery characterized by increased IL-10, TNF-α and SA-β-gal, reduced incidence of apoptosis and altered metabolism. Our findings suggest that this senescent-like phenotype is associated with immature PMNs that are elevated in aging and that these cells are more likely to display this senescent-like phenotype. It is possible that senolytic or senomorphic therapeutics, which clear senescent cells or inhibit production of SASP factors (L. Zhang, Pitcher, Prahalad, Niedernhofer, & Robbins, 2023), may rescue PMN anti-pneumococcal responses in old hosts. These therapies have been shown to improve immune responses following treatment of senescent T cells, B cells, macrophages, and hematopoietic cells in malignancies (Cai et al., 2020; Chang et al., 2016; Cobanoglu et al., 2023; Yousefzadeh et al., 2021). Recent studies have also found removal of senescent-like PMNs to be beneficial in tumor models (Bancaro et al., 2023; Guo et al., 2023). In infection, studies in hamsters and mice have shown that senolytics significantly decreased serious illness and improved survival upon SARS-CoV 2 infection, although the effect on PMNs was not directly tested (Camell et al., 2021; X. Zhang, Suda, & Zhu, 2023). In summary, this study provides evidence for an age associated senescence-phenotype in PMNs that is context dependent suggesting that strategies targeting senescent-like PMNs could improve disease outcomes in the future.

## Materials and Methods

### Animals

Young (8–12 weeks) and old (20–22 months) C57BL/6 male mice were obtained from the National Institute on Aging colony or from Jackson Laboratories (Bar Harbor, ME). Experiments were performed in males as they are more susceptible to pneumococcal infection (Gutierrez et al., 2006). All mice were placed at the University at Buffalo in specific pathogen free housing for at least 2 weeks prior to use. Ets1KO mice were generated as previously described (Barton et al., 1998) and maintained on a mixed 129/SVxC57BL/6 background (D. Wang et al., 2005). All experiments were conducted in accordance with the Institutional Animal Care and Use Committee guidelines.

### Human Donors

Male and female donors, 24-41 years old, were recruited. Individuals taking medications, pregnant, or with infections within the last two weeks were excluded. All enrolled donors signed approved informed consent forms in conjunction with IRB approval. Blood was collected into tubes containing sodium citrate at 9AM to control for circadian rhythm.

### Bacteria

*Streptococcus pneumoniae* TIGR4 strain was a kind gift from Andrew Camilli. Briefly, bacteria were grown to mid-exponential phase at 37°C at 5% CO_2_ in Todd Hewitt broth supplemented with 0.5% yeast extract and Oxyrase as previously described (Siwapornchai et al., 2020). Heat-killed (HK) *Sp* was generated as described (Herring et al., 2022), by incubation at 55°C for 2 hours.

### Mouse Infections

Anesthetized mice were infected by assisted aspiration using the tongue pull method that we call intratracheal (i.t.) with 1×10^6^ CFU of *Sp* where the bacterial inoculum is pipetted into the trachea with the tongue gently pulled out to ensure direct delivery into the lungs. Animals were euthanized for organ harvest.

### PMN Isolation

For RNAseq: PMN negative selection kit (customized; StemCell) was used per the manufacturer’s protocol to obtain a highly enriched population. Briefly, lungs were harvested at 12- and 24HPI, digested as described (Herring et al., 2022) and used for PMN isolation. PMN purity was determined by flow cytometry (**SF1B**).

For opsonophagocytic killing assays: Bone Marrow (BM) cells were collected from the femurs and tibias of uninfected mice via density-gradient centrifugation using histopaques as previously described (Bhalla et al., 2020).

For Seahorse Metabolic Assays: BM cells were collected from the femurs and tibias of uninfected mice. Cells were flushed and red blood cells were then lysed using ACK lysis buffer, and remaining pellet was resuspended in HBSS. PMNs were then isolated using the StemCell EasySep Mouse Neutrophil Enrichment Kit.

For Human peripheral blood PMNs: PMNs were isolated utilizing the StemCell EasyStep Human PMN Isolation Kit as per manufacturer’s protocol.

### RNA purification, Illumina library preparation, and RNA Sequencing Analysis

RNA was isolated from PMNs using a RNeasy Mini Kit according to the manufacturer’s specifications. RNA quality, cDNA library prep, sequencing, and data processing was conducted as previously described (Bhalla et al., 2021). We defined significant up- or downregulation of gene expression as an absolute fold change of ≥.75 with an p-adj value of <.05. Heatmaps were using normalized transcript per million (TPM) values on the webtool heatmapper.ca (Babicki et al., 2016).

### Pathway Analysis

Pathway and functional enrichment analysis were conducted using the Database for Annotation, Visualization, and Integrated Discovery (DAVID). Terms and pathways with a p value<.05 were considered significant. For pathways of the Kyoto Encyclopedia of Genes and Genomes (KEGG), the pathway was retrieved from https://www.genome.jp/kegg/ and edited using GNU Image Manipulation Program to colorize differential expression.

### Transcription Factor Signature Enrichment and Protein Interaction Network Analysis

DEGs identified were input into the ChEA3 online webtool and analyzed using TopRanked scoring (Keenan et al., 2019). TFs with a score <.01 were considered significantly enriched. TFs identified as significantly enriched were input into the STRING interaction network analysis webtool (Szklarczyk et al., 2023). The interaction score was set to high confidence (0.7). Clusters were identified using the MCL clustering (inflation parameter =2.0). The resulting clustered interaction network was edited using GIMP to colorize to indicate enrichment in young (yellow), old (pink), or both (black). TRANSFAC analysis was conducted using gProfiler functional enrichment analysis webtool utilizing default settings (Kolberg et al., 2023).

### Hoxb8 PMN Differentiation

Hoxb8 cell lines were maintained and differentiated as previously described (Nguyen et al., 2020). In brief, Hoxb8 progenitors were maintained in RPMI 1640 supplemented with 10% FBS, β-estradiol (E2), and stem cell factor (SCF). Cells were washed three times in PBS and cultured in E2-free RPMI 1640 supplemented with FBS, G-CSF, IL-3, and SCF for 2 days. Cells were then resuspended in E2-free RPMI supplemented with FBS and G-CSF for one day.

### Extracellular Flux Assays

Extracellular flux assay protocols were modified from previous protocols (Grudzinska et al., 2023; Mathuram et al., 2022). In brief, PMNs resuspended in RPMI1640 supplemented with glucose, pyruvate, glutamine, and FBS were seeded on Seahorse Cell Culture plate precoated with murine sera. Extracellular flux was measured using a Seahorse Xfe96 analyzer using the setups below.

For respiration: Oxygen consumption rate was obtained following sequential treatment with 2.5uM oligomycin, FCCP, and 1uM mixture of rotenone/antimycin A. Basal respiration, max respiration, respiratory capacity, proton leak, and ATP production were calculated as previously described (Mathuram et al., 2022).

For glycolysis: Extracellular acidification rate (ECAR) was converted to proton efflux rate (PER) utilizing Agilent Wave software following treatment with Vehicle Control (VC) (unbuffered RPMI1640), murine sera opsonized Heat-killed *Sp* (MOI=10), and 60mM 2-deoxyglucose (2-DoG). Basal glycolysis and max glycolysis were calculated as described previously (Grudzinska et al., 2023).

### Opsonophagocytic Killing Assay

The ability of PMNs to kill *S. pneumoniae ex vivo* was measured using an opsonophagocytic killing assay as previously described (Bou Ghanem et al., 2015). Briefly, PMNs treated with VC (DMSO), lonidamine, 2-DoG at the indicated concentrations were infected with *Sp* (MOI=.01) opsonized with mouse serum. Reactions were rotated at 37°C for 45 min. Percent killing was determined plating on blood agar plates in comparison to the no-PMN controls.

### RT-qPCR

RNA was isolated from the indicated samples utilizing a Qiagen RNeasy Mini Kit as per the manufacturer’s instructions. cDNA conversion was done using the SuperScript VILO cDNA Synthesis kit. RT-qPCR was done using iQ SYBR Green Supermix to detect relative levels of transcript cDNA using the following primers in **Table S2**

### Flow Cytometry

The lungs were digested as previously described (Siwapornchai et al., 2020). Splenocytes were harvested following physical disruption of the spleen. BM cells were collected from the femurs and tibias. Red blood cells were lysed using ACK lysis buffer, and remaining pellet was resuspended in HBSS. Cells were stained with antibodies, kits, and stains purchased from BD Biosciences, Invitrogen, Biolegend, and Dojindo Molecular Technologies used as listed in **Table S3**. 20,000-50,000 events were collected utilizing a BD Celesta or BD Fortessa Cytometer. Data were analyzed using FlowJo with representative gating depicted in **SF18**.

### Statistics

Data was analyzed using GraphPad Prism 10. Data are presented as mean and standard deviation. All data were tested for normality using Shapiro-Wilk Test. Significance was tested using Student’s t test, Mann-Whitney test, one-sample t test (different that 1), One-way-ANOVA followed by Tukey’s multiple comparisons test or Kruskal-Wallis test as appropriate (ns not significant, *p<.05, ** p<.01, *** p<.001, **** p<.0001).

## Author Contributions

MCB and MB conducted research, analyzed data, and wrote paper. BM, AB, APL, RH, MCK, LH conducted research. PAL, LAGS, JM, and ABP provided materials. ENBG designed research, edited the paper, and had responsibility for final content. All authors read and approved the manuscript.

## Data Availability

RNA-Seq Data are publicly available on Gene Expression Omnibus (GEO) with accession number GSE294007. All other data are available from the corresponding author upon request.

