## Supplemental Data for "Host aging induces a senescent-like phenotype in neutrophils and altered transcriptional responses to *Streptococcus pneumoniae*"

### Supplemental Materials

### Supplemental Figure Legends

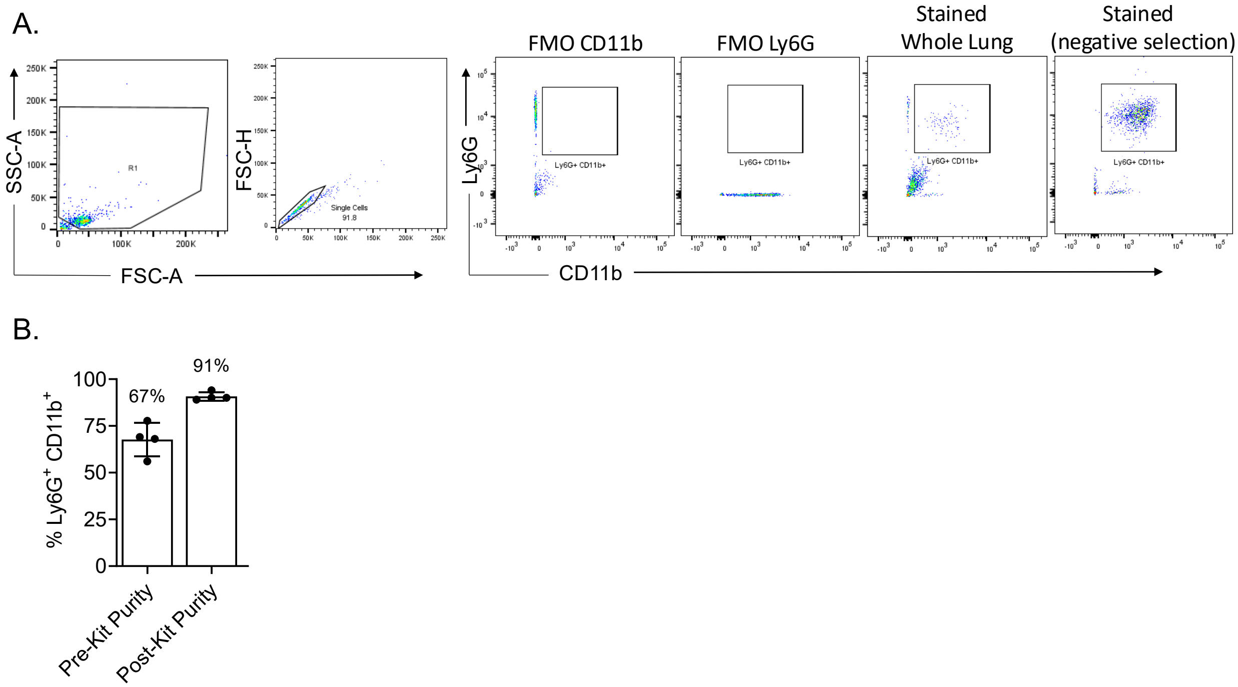


**SF1. Verification of purity of PMNs sent for RNA sequencing.** Gating strategy utilized to confirm purity of pre and post-enrichment for PMNs analyzed via RNAseq (A). Observed purity of samples pre-enrichment and post-enrichment utilizing a customized StemCell negative selection purification kit with each point representing a biological replicate (B).

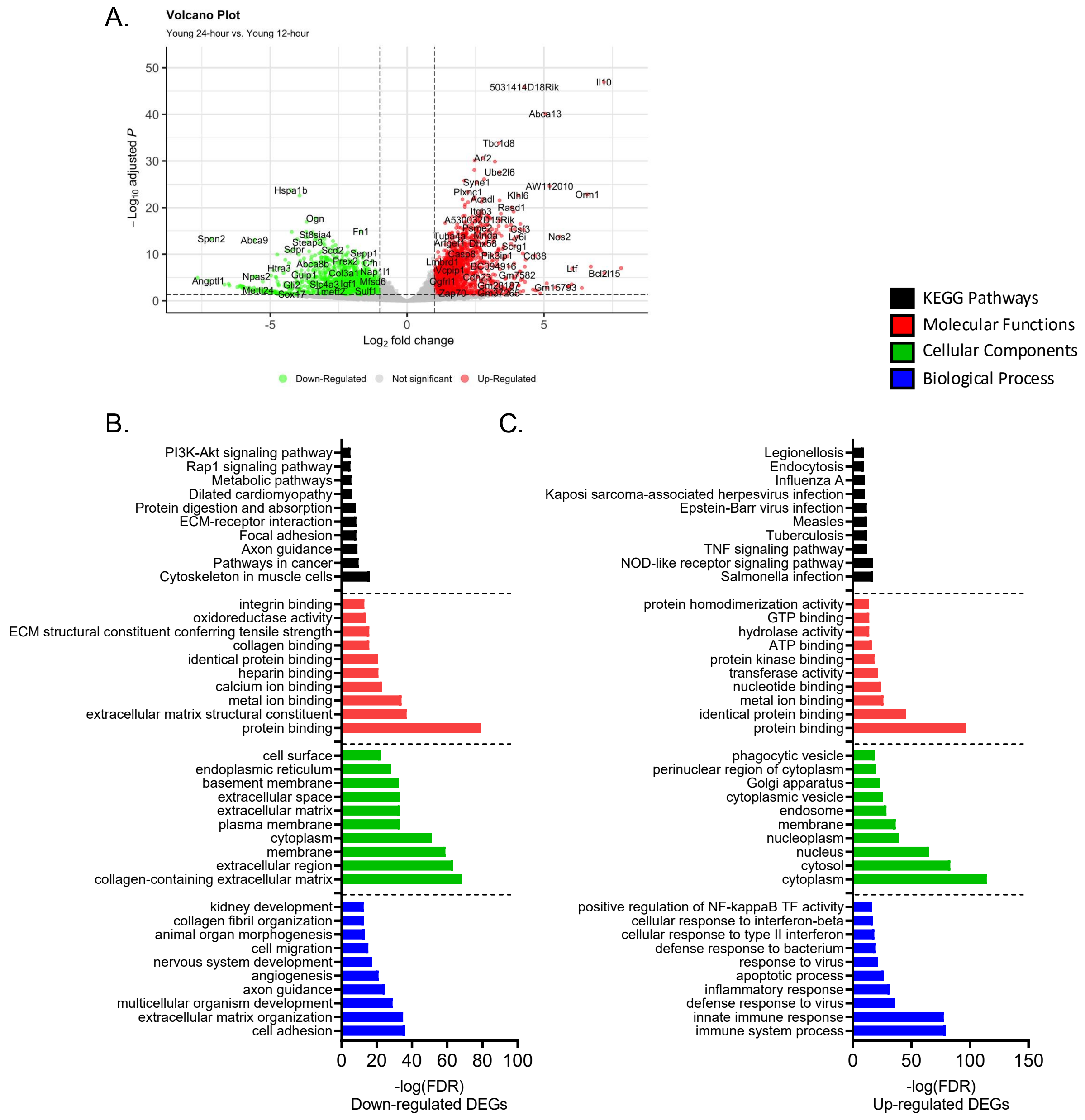


**SF2. Gene expression changes in PMNs isolated from young hosts are associated with normal inflammatory response to *S. pneumoniae*.** Volcano plot indicating DEGs identified between Young-24HPI and Young-12HPI (A). Top 10 results of DAVID analysis results for terms enriched in DEGs identified as downregulated in Young-24HPI vs Young-12HPI (B). Top

10 results of DAVID analysis results for terms enriched in DEGs identified as upregulated in Young-24HPI vs Young-12HPI (C).

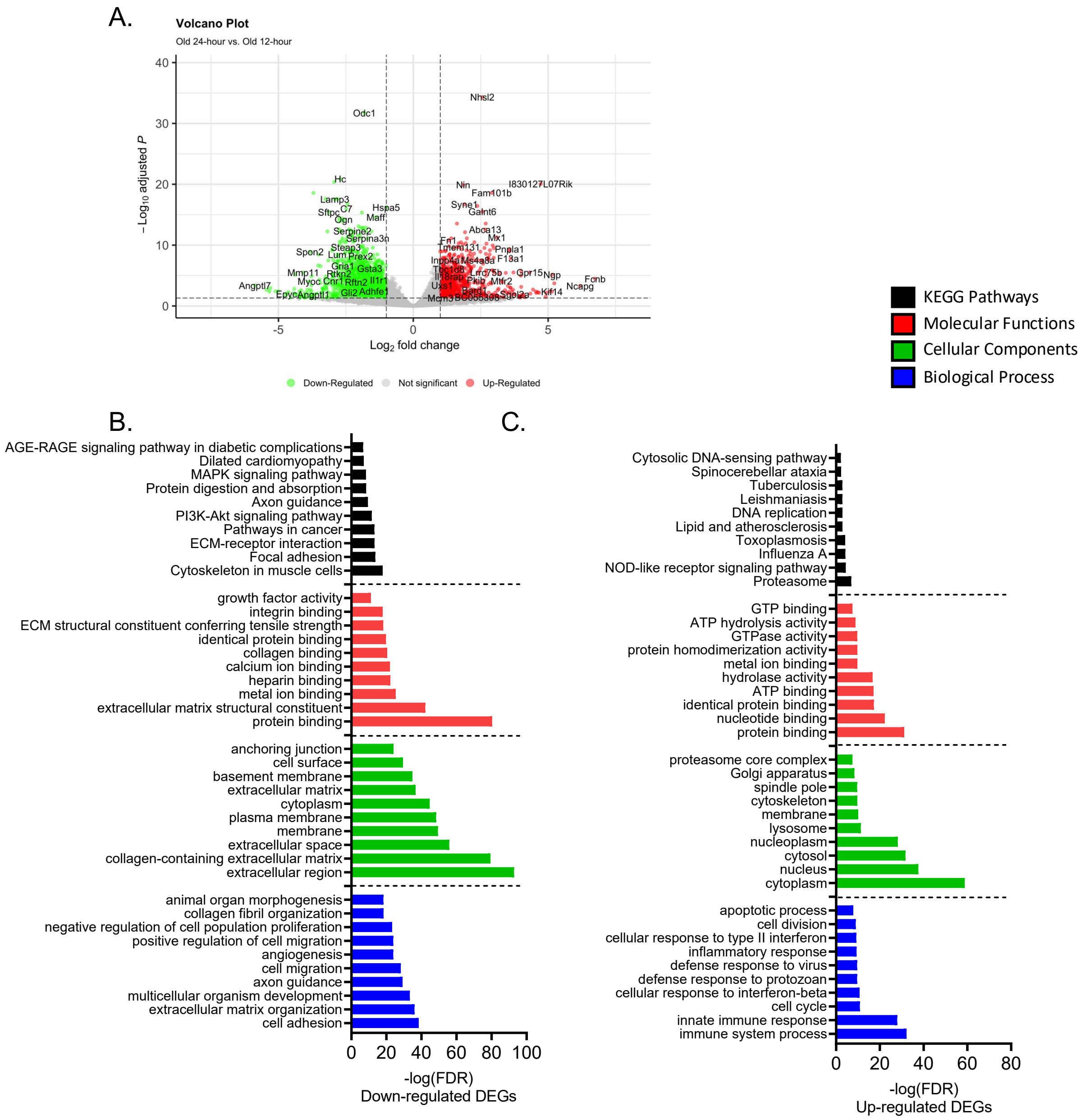


**SF3. Gene expression changes in PMNs isolated from aged hosts are associated with a muted inflammatory response to *S. pneumoniae*.** Volcano plot indicating DEGs identified between Old-24HPI and Old-12HPI (A). Top 10 results of DAVID analysis results for terms enriched in DEGs identified as downregulated in Old-24HPI vs Old-12HPI (B). Top 10 results of

DAVID analysis results for terms enriched in DEGs identified as upregulated in Old-24HPI vs Old-12HPI (C).

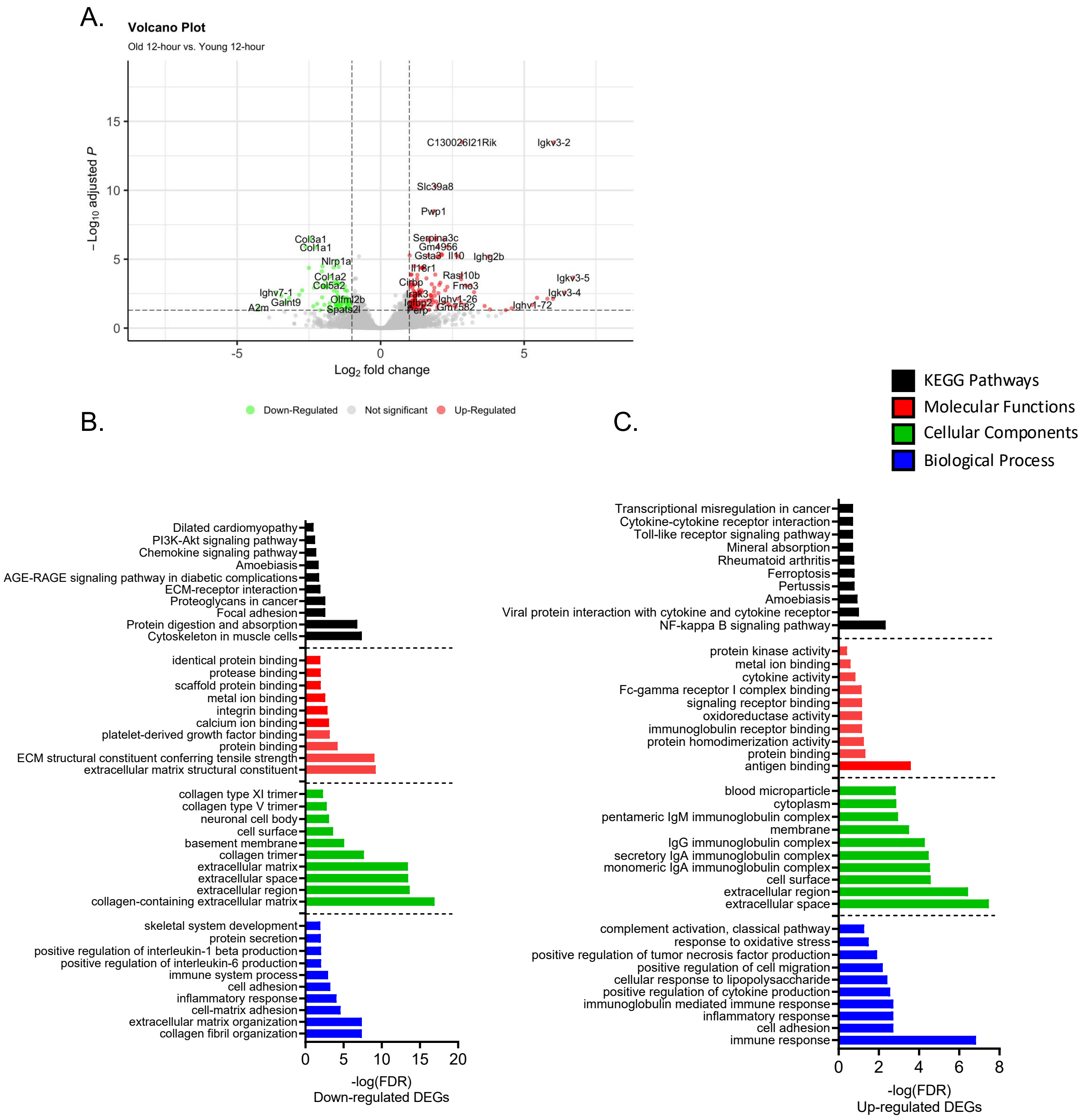


**SF4. Transcriptome of lung PMNs 12HPIs are similar across host age*.*** Volcano plot indicating DEGs identified between Old-12HPI and young-12HPI (A). Top 10 results of DAVID analysis results for terms enriched in DEGs identified as downregulated in Old-12HPI vs Young-12HPI (B). Top 10 results of DAVID analysis results for terms enriched in DEGs identified as upregulated in Old-12HPI vs Young-12HPI (C).

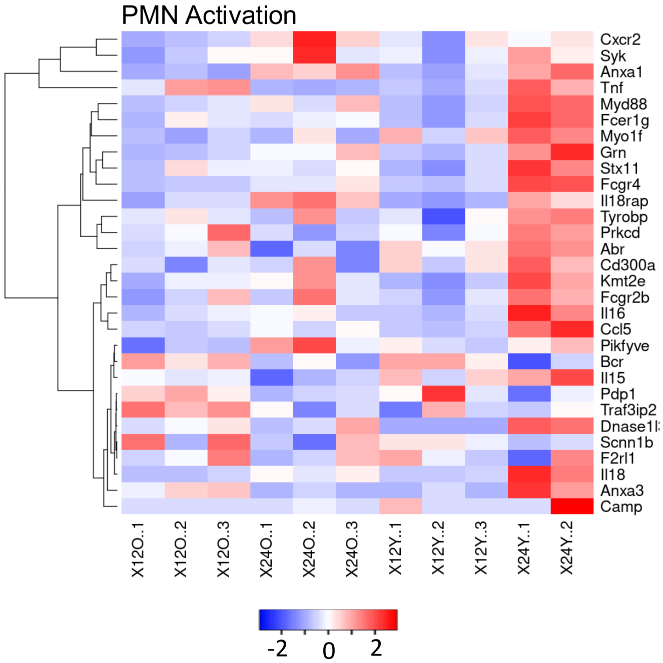


**SF5. Aging is associated with decreased expression for genes associated with PMN activation.** Heatmap showing normalized gene expression for genes associated with the GO terms Neutrophil Activation, Positive Regulation of Neutrophil Activation, and Negative Regulation of Neutrophil Activation.

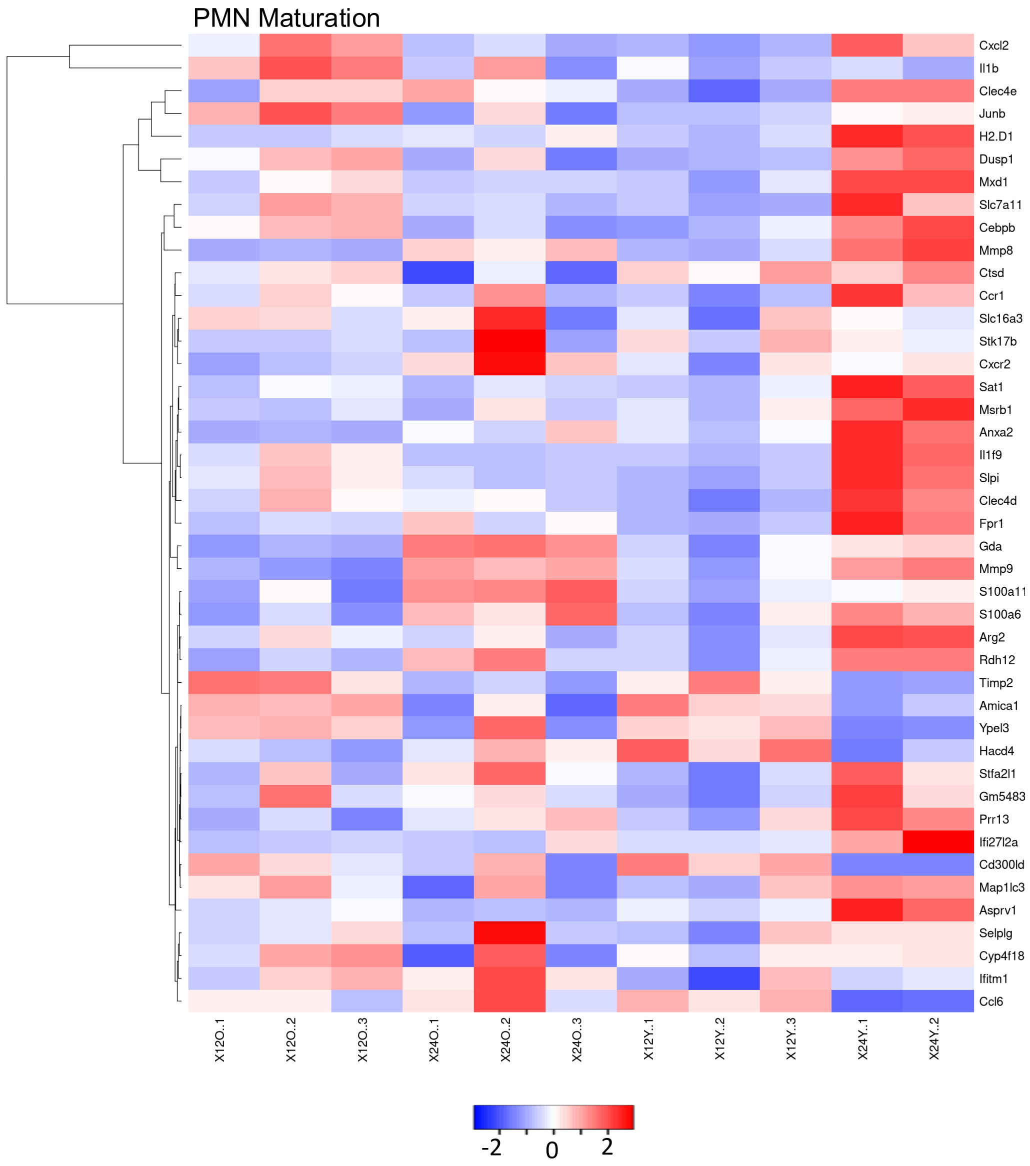


**SF6. Aging is associated with decreased expression for genes associated with PMN maturation.** Heatmap showing normalized gene expression for genes identified by Ai et al 2022.

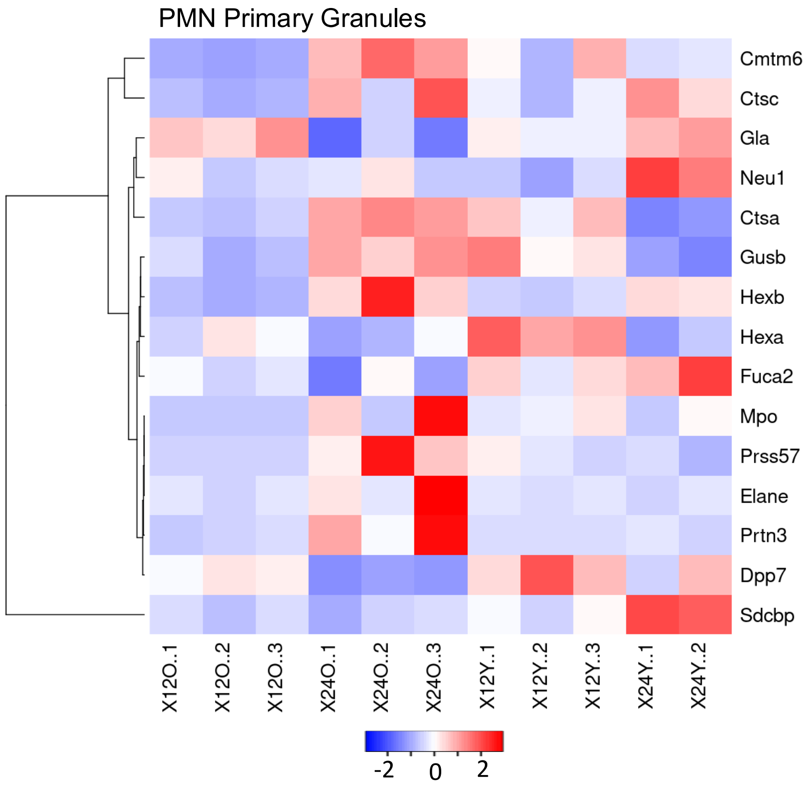


**SF7. Aging is associated with altered expression of primary granule contents.** Heatmap showing normalized expression of genes associated with primary granules.

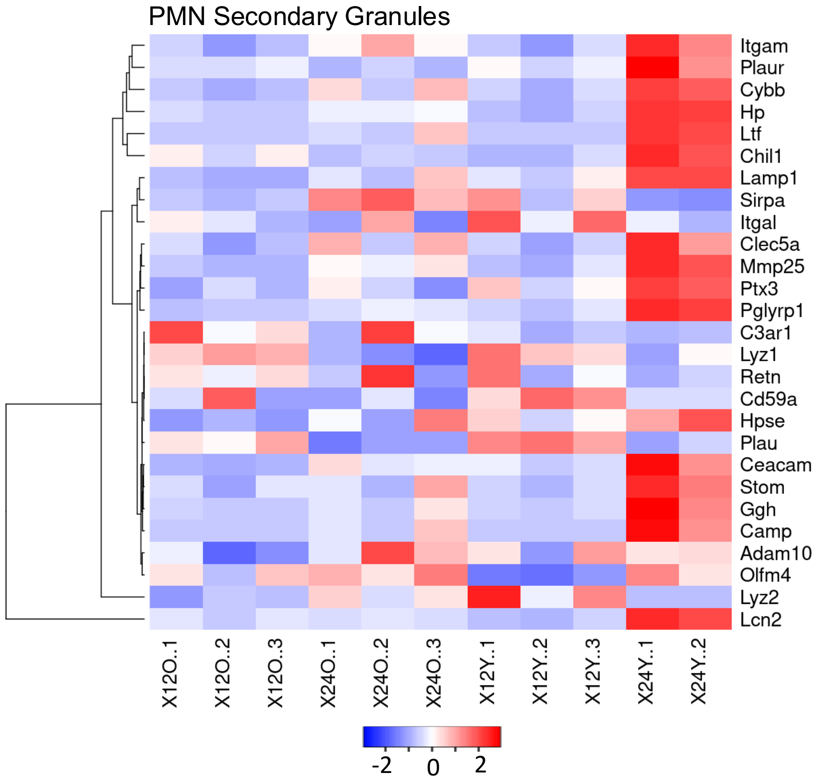


**SF8. Aging is associated with altered expression of secondary granule contents.**

Heatmap showing normalized expression of genes associated with secondary granules.

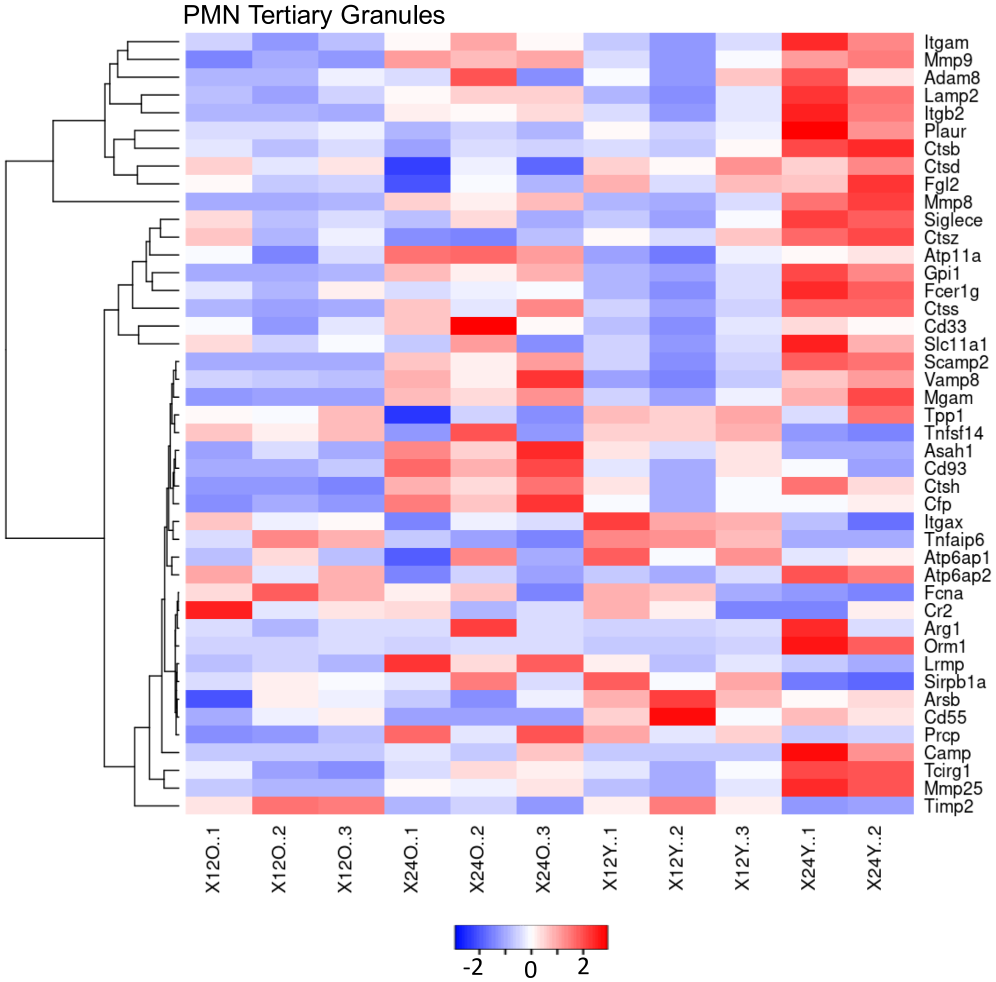


**SF9. Aging is associated with altered expression of tertiary granule contents.** Heatmap showing normalized expression of genes associated with tertiary granules.

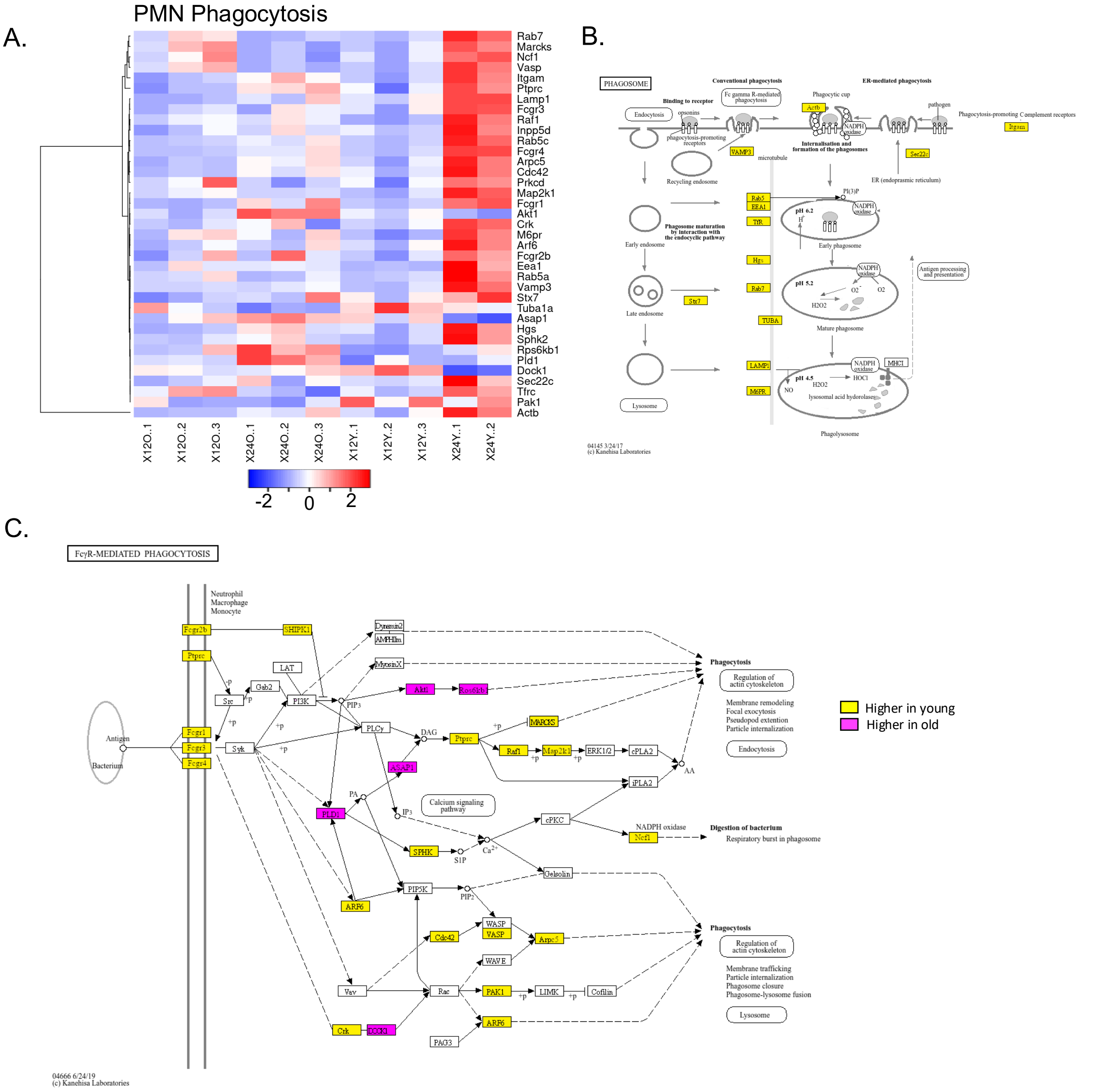


**SF10. Aging is associated with impaired upregulation of genes involved in phagocytosis.** Heatmap showing normalized expression of genes part of the Fc-gamma Mediated Phagocytosis and Phagosome KEGG pathways (A). The KEGG pathway Phagosome pseudocolored to indicate higher expression of genes in PMNs isolated from young (yellow) or old (pink), or neither (white) (B). The KEGG pathway Fc-gamma Mediated Phagocytosis

pseudocolored to indicate higher expression of genes in PMNs isolated from young (yellow), old (pink), or neither (white) (C).

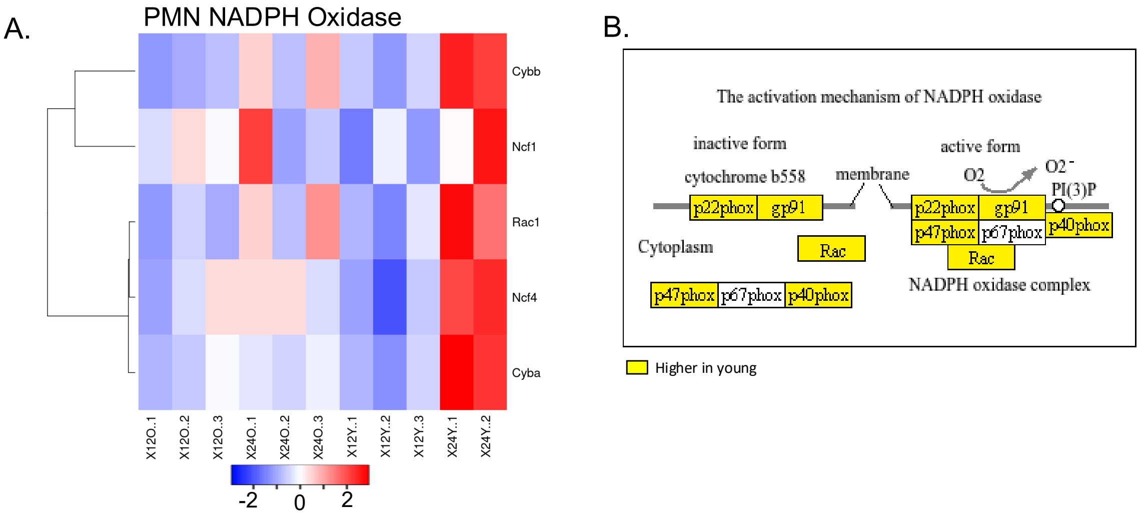


**SF11. Aging is associated with impaired upregulation of genes involved in NADPH oxidase complex.** Heatmap showing normalized expression of genes associated with the activation mechanism of NADPH oxidase section of the KEGG pathway Phagosome (A). The activation mechanism of NADPH oxidase region of the KEGG pathway Phagosome pseudocolored to indicate higher expression of genes in PMNs isolated from young (yellow), old (pink), or neither (white) (B).

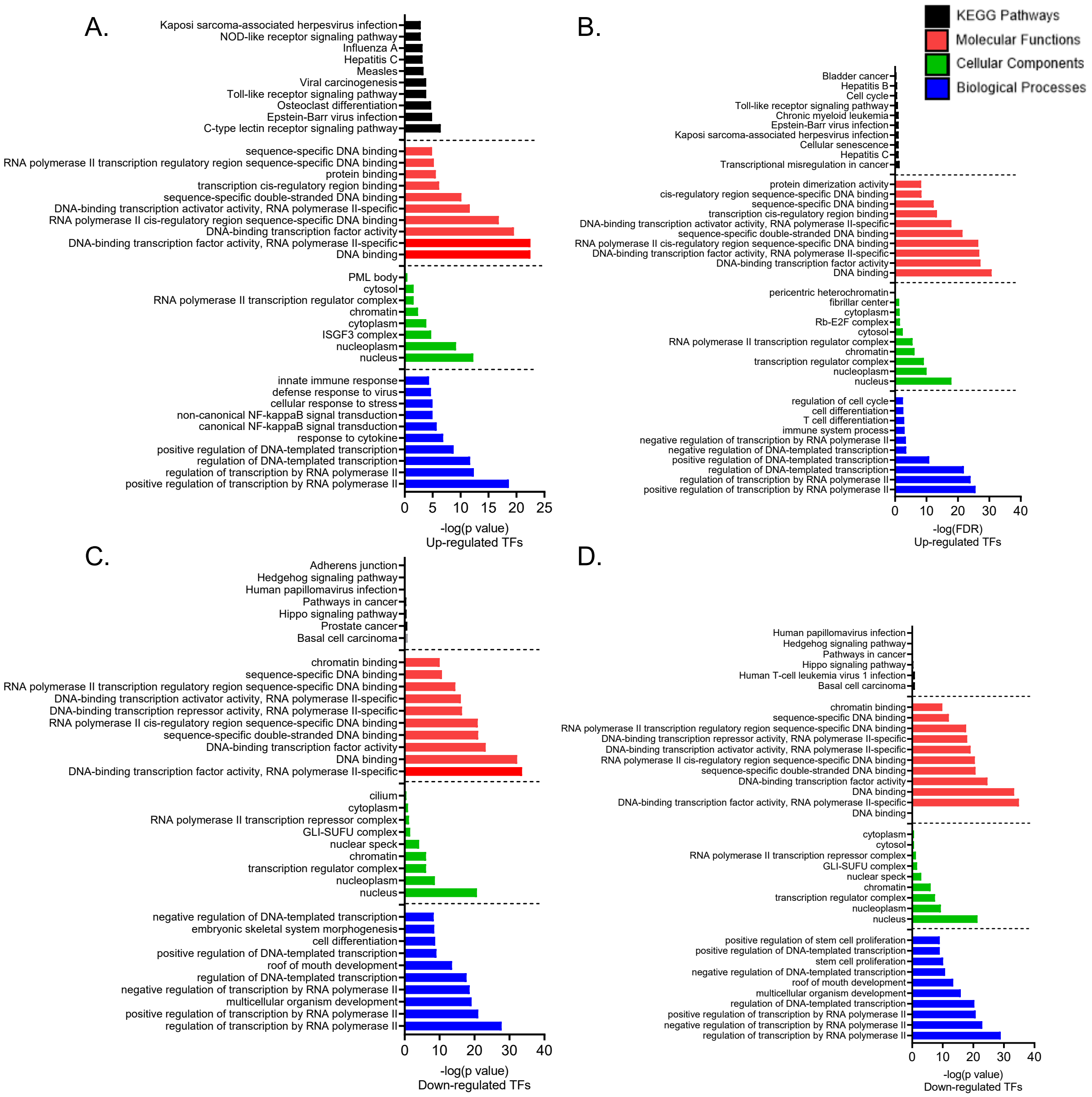


**SF12. Significantly enriched transcription factors in young versus old mice display different pathway involvement.** Top 10 results for DAVID analysis conducted on transcription factors identified as significantly enriched by ChEA3 in DEGs identified as upregulated in PMNs isolated from the lungs of young mice (A). Top 10 results for DAVID analysis conducted on transcription factors identified as significantly enriched by ChEA3 in DEGs identified as

upregulated in PMNs isolated from the lungs of aged mice (B). Top 10 results for DAVID analysis conducted on transcription factors identified as significantly enriched by ChEA3 in DEGs identified as downregulated in PMNs isolated from the lungs of young mice (C). Top 10 results for DAVID analysis conducted on transcription factors identified as significantly enriched by ChEA3 in DEGs identified as downregulated in PMNs isolated from the lungs of aged mice (D).

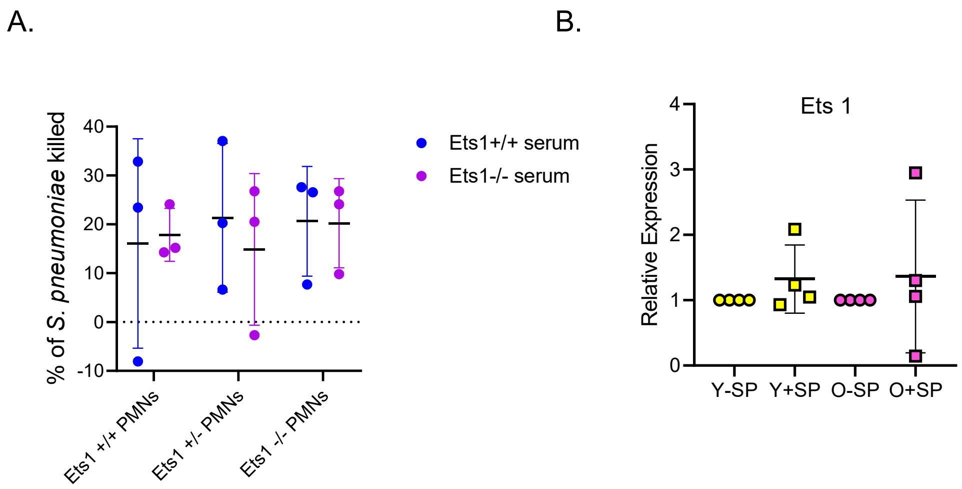


**SF13. Ets1 is not involved in PMN responses to *S. pneumoniae*.** Representative results from one of three separate experiments of *S. pneumoniae* opsonophagocytic killing assays conducted using PMNs isolated from the BM of Ets1 +/+, Ets1 +/-, and Ets1-/- mice using sera from Ets1+/+ and Ets1-/- as an opsonin. Each point represents a technical replicate (A). Summary RT-qPCR data showing Ets1 relative mRNA transcript levels in human peripheral blood PMNs mock infected or infected with *S. pneumoniae* (MOI 2). Each point represents a biological replicate (B).

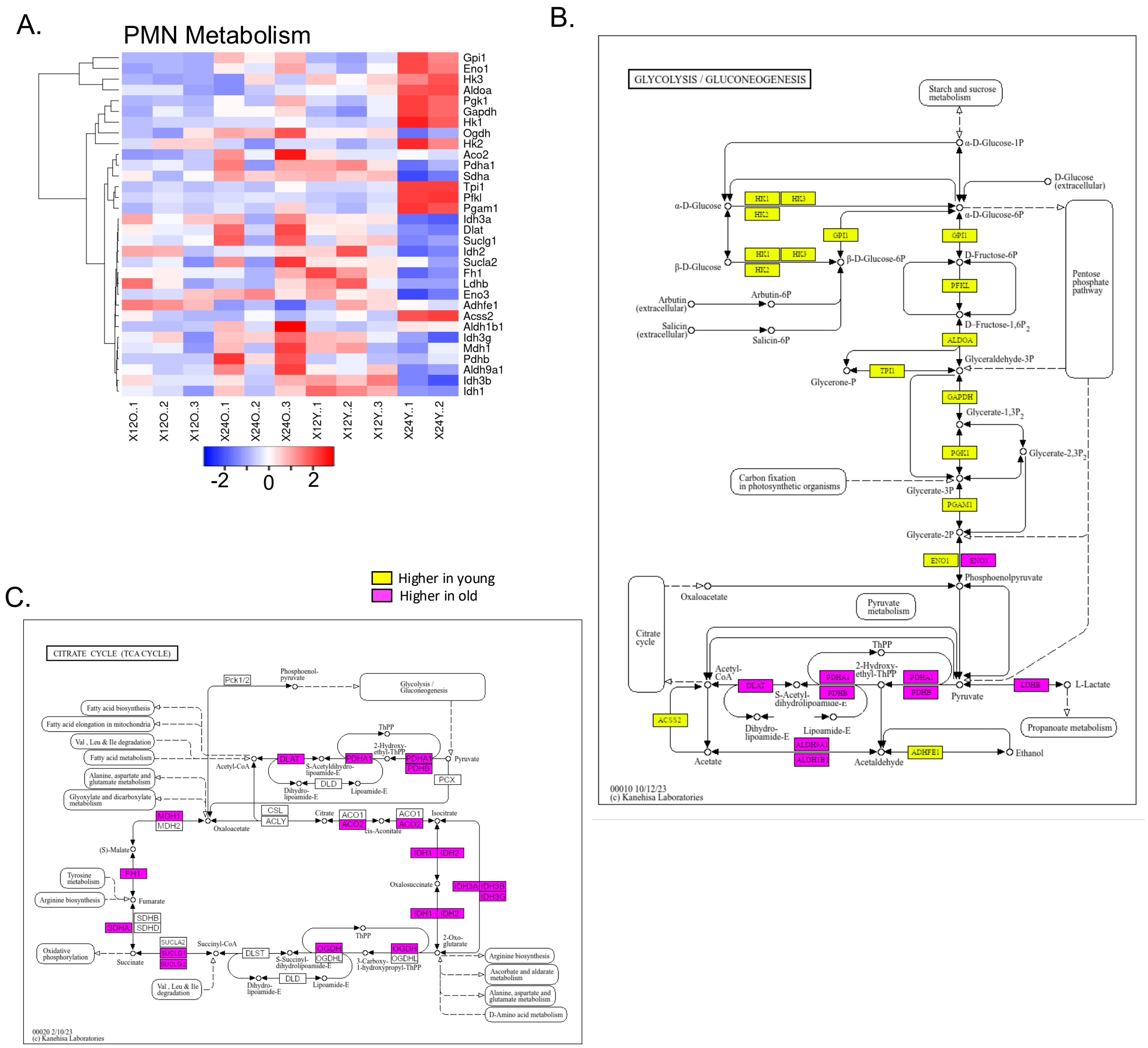


**SF14. Aging is associated with significant changes in metabolic responses to *S. pneumoniae* infection in lung PMNs.** Heatmap showing normalized expression of genes associated with the KEGG pathways Glycolysis/Gluconeogenesis and Citrate Cycle (TCA Cycle) (A). The KEGG pathway Glycolysis/Gluconeogenesis pseudocolored to indicate higher expression of genes in PMNs isolated from young (yellow), old (pink), or neither (white) (B). The KEGG pathway Citrate Cycle (TCA Cycle) pseudocolored to indicate higher expression of genes in PMNs isolated from young (yellow), old (pink), or neither (white) (C).

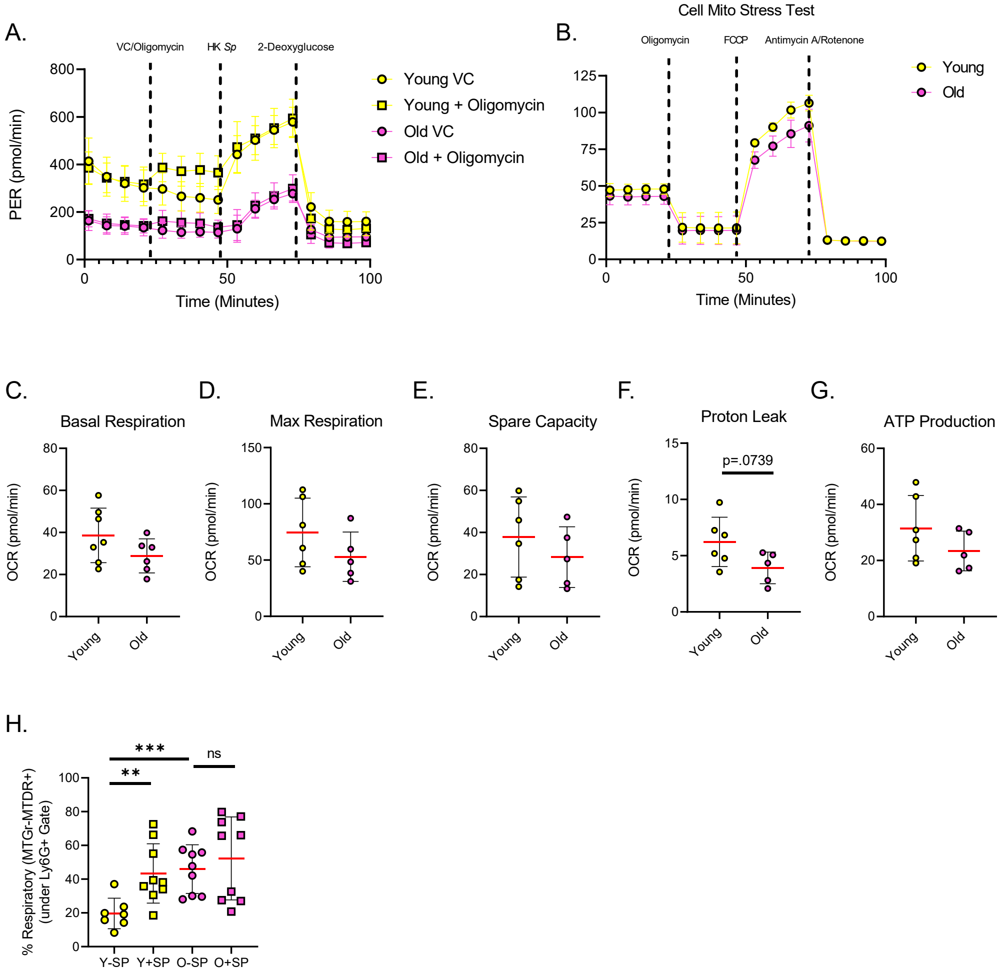


**SF15. Aging is not associated with significantly differential respiration in bone marrow derived PMNs.** Representative results of Glycolysis Stress Test conducted on BM PMNs from young and aged mice in the presence of the respiration inhibitor oligomycin (A). Representative results of Cell Mito Stress test conducted on BM PMNs isolated from young and aged mice (B). Summary data showing basal respiration (C), max respiration (D), spare capacity (E), proton leak (F), and ATP production (G) measurements as derived from Cell Mito Stress Test. Summary data from flow cytometric analysis of lung PMNs showing % respiratory (Mitotracker

Green-Mitotracker Deep Red+). For C-H data are pooled from three separate experiments and each point represents a biological replicate.

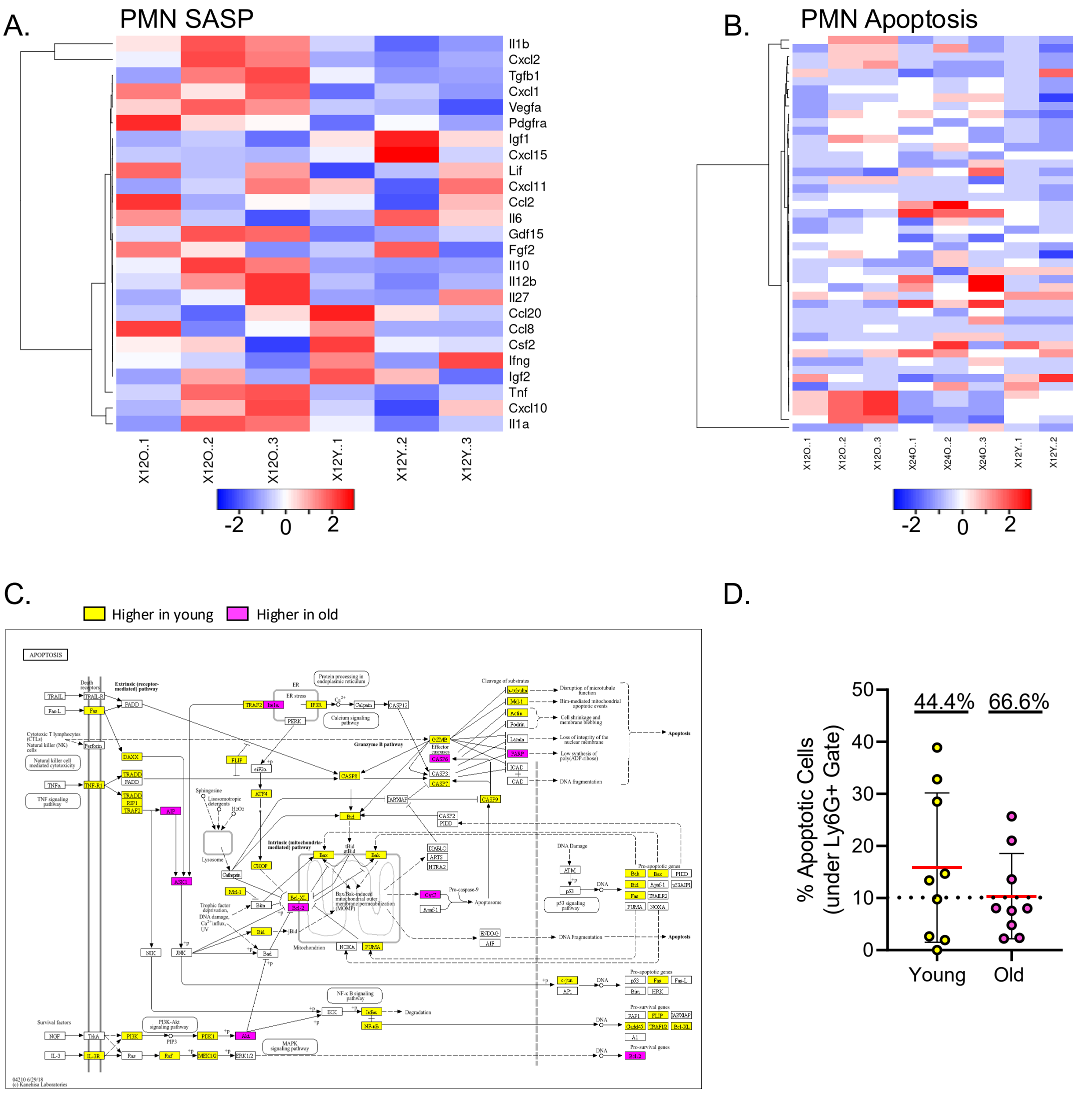


**SF16. Aging is associated with altered secretory factor expression and apoptosis induction.** Heatmaps showing normalized expression of SASP factors (A) and genes associated with the KEGG pathway Apoptosis (B). The KEGG pathway Apoptosis pseudocolored to indicated elevated expression in young (yellow), aged (pink), or neither (white) 24HPI (C). Flow cytometry data showing % apoptotic PMNs in the lungs of young and old mice highlighting the percentage of mice displaying less than 10% apoptotic PMNs (D). Data are

pooled from three separate experiments and each point represents a biological replicate. Significance was determined using Mann-Whitney test.

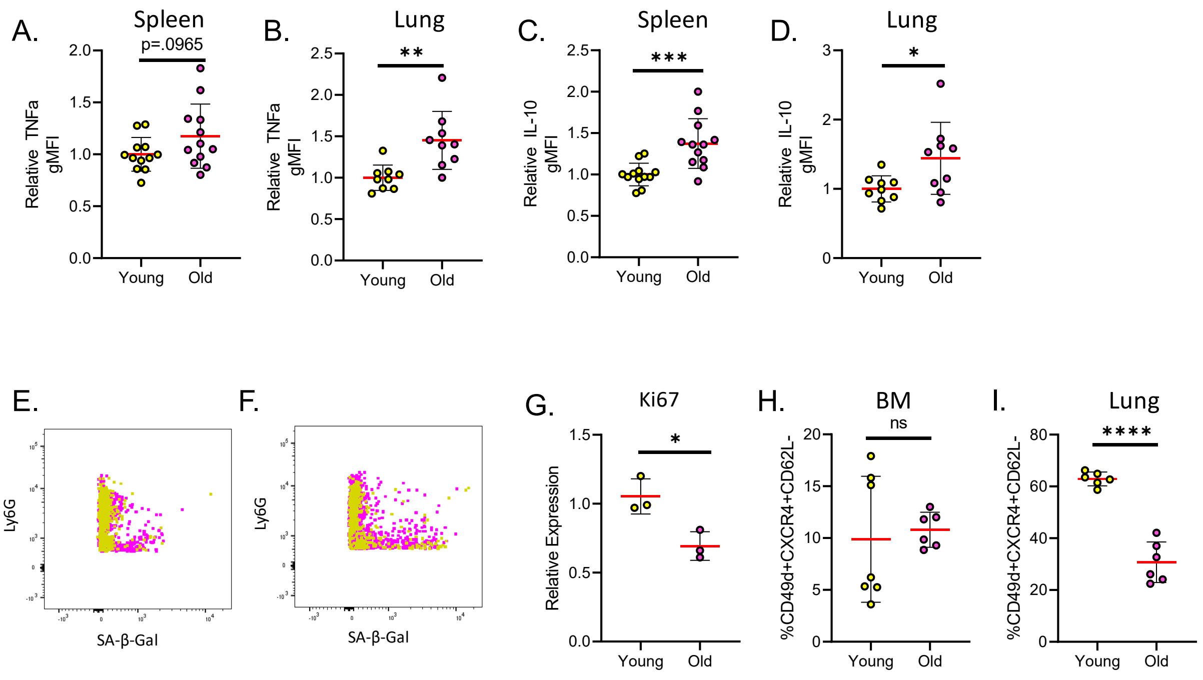


**SF17. Aging is associated with a senescent-like phenotype in PMNs.** Data showing TNFα (A) and IL-10 (B) gMFI of PMNs within the spleen from young and aged mice relative to average of young controls. Data showing TNFα (A) and IL-10 (B) gMFI of PMNs within the lung from young and aged mice relative to average of young controls. Representative flow plots of SA-β-Gal vs Ly6G in PMNs from young (yellow) and old (pink) mice in the spleen (E) and lungs (F). Data showing relative transcript levels for Ki-67 in BM PMNs isolated from young and old mice (G). Flow cytometry data for % of Ly6G+ cells that are CD49d+CXCR4+CD62L in the BM (H) and lung (I). Each point represents a biological replicate pooled from three separate experiments. Significance was determined using Unpaired t-tests or Mann-Whitney test following testing of normality by Shapiro-Wilk Test.

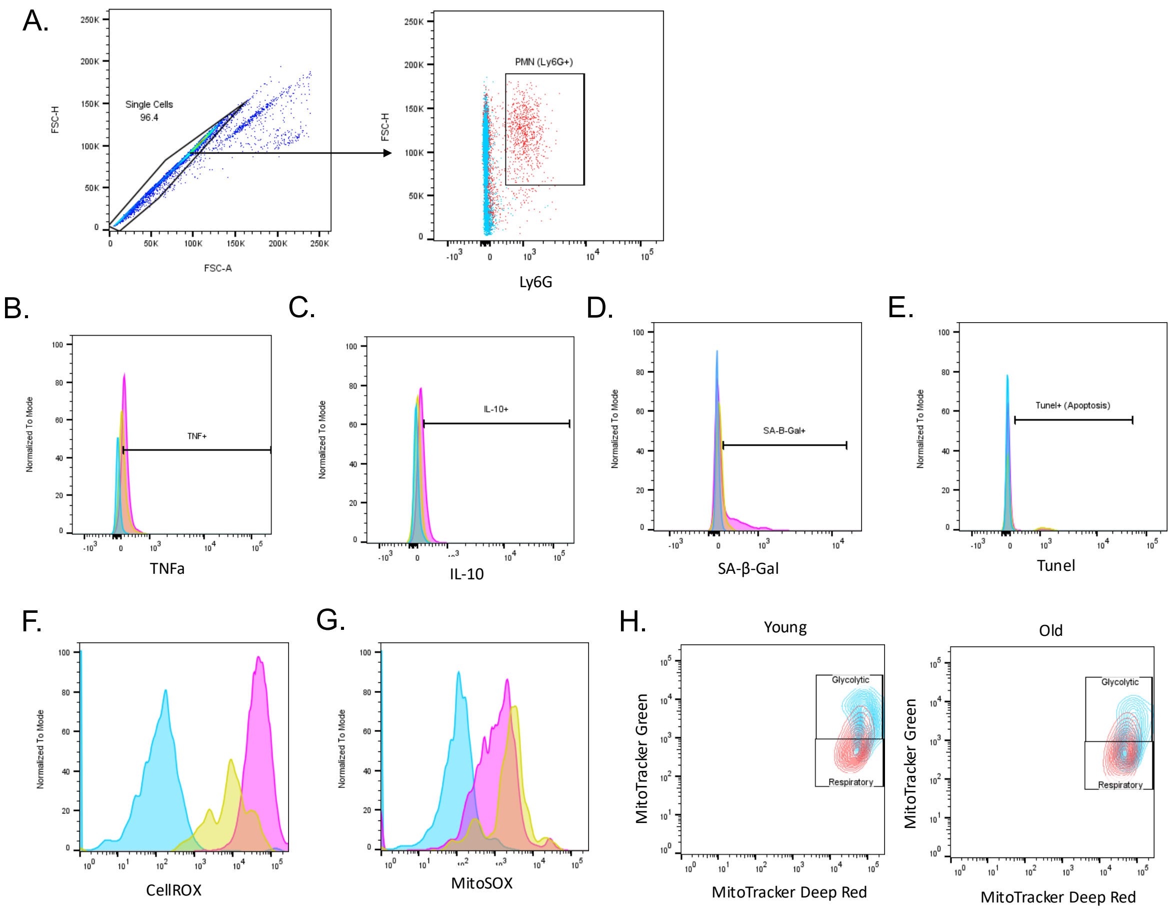


**SF18. Representative gating strategy used for flow cytometry analysis of PMNs *in vivo.***

Representative gating used to define singlets and Ly6G+ PMNs (Blue=Ly6G FMO, Red=Stained) (A). Representative flow plots and histograms depicting staining for IL-10 (B), TNFα (C), SA-β-Gal (D), Tunel (apoptosis), CellROX (F), and MitoSOX (G) on PMNs as defined above (Blue= FMO, Yellow=Stained Young, Pink=Stained Old). Representative flow plots depicting gating strategy for determination of glycolytic (MitoTracker Green+ MitoTracker Deep Red+) and respiratory (MitoTracker Green- MitoTracker Deep Red+) PMNs in young and old mice displaying uninfected groups in blue and infected groups in red (H).

| Color | ID | Description |
|---|---|---|
| | 1 | Interferon Regulatory Factor |
| | 2 | Mixed: C-terminal binding protein, drought induced 19 protein type, zinc-binding |
| | 3 | Negative regulation of smoothened signaling pathway |
| | 4 | Interferon-regulatory factor 3 |
| | 5 | Erythroblast transformation specific domain |
| | 6 | Regulation of transcription in G1/S transition of mitotic cell cycle |
| | 7 | Ig-like, plexins, transcription factors |
| | 8 | Myogenesis |
| | 9 | Domain first found in mice T locus (Brachyury) protein |
| | 10 | Mixed: Vitellogenesis and F-actin binding |
| | 11 | Middle Ear Morphogenesis |
| | 12 | Myeloid dendritic cell differentiation |
| | 13 | Mixed: Primitive hemopoiesis and Helix-loop-helix protein TAL-like |

**Supplemental Table 1. ChEA3 STRING clusters.** Interaction network clusters as determined by STRING analysis of transcription factors identified by ChEA3 analysis of up- and down-regulated DEGs in both young and old mice.

| Target | Forward Primer | Reverse Primer |
|---|---|---|
| **h-Ets1** | TCAAACAAGAAGTCG TCACC | AAGCTGTCATAGGAGGGAAC) |
| **h-GAPDH** | AGCCACATCGCTCAGACAC | GCCCAATACGACCAAATCC |
| **m-FasL** | TGGGTAGACAGCAGTGCCAC | GCCCACAAGATGGACAGGG |
| **m-p16** | GGGCTCGGCTGGATGTG | CTTGATGTCCCCGCTCTTGG |
| **m-Actin** | GCAGCTCCTTCGTTGCCGGTC | TTTGCACATGCCGGAGCCGTTG |

**Supplemental Table 2. qPCR Primers.** Targets and primer sequences used for qPCR analysis of human and murine samples.

| Stain | Clone or Catalogue # |
|---|---|
| **Fc-Block** | 2.4G2 |
| **Anti-Ly6G** | 1A8 |
| **anti-CD11b** | M1/70 |
| **anti-IL-10** | JES5-16E3 |
| **anti-TNF** | MP6-XT22 |
| **Anti-CD49d** | R1-2 |
| **Anti-CD62L** | MEL-14 |
| **Anti-CXCR4** | 2B11 |
| **SA-β-gal** | Cat # NC1610927 |
| **MitoSOX Red** | Cat # M36008 |
| **CellROX Deep Red** | Cat # C10422 |
| **Coralite504Tunel** | PF00009 |
| **MitoTracker Green FM** | M46750 |
| **MitoTracker Deep Red FM** | M46753 |

**Supplemental Table 3. Flow cytometry reagents.** Antibodies and kits utilized for staining and flow cytometry analysis.
